# An evolutionary perspective on the origin, conservation and binding partner acquisition of tankyrases

**DOI:** 10.1101/2022.04.24.489311

**Authors:** Sven T. Sowa, Chiara Bosetti, Albert Galera-Prat, Mark S. Johnson, Lari Lehtiö

## Abstract

Tankyrases are poly-ADP-ribosyltransferases that regulate many crucial and diverse cellular processes in humans such as Wnt signaling, telomere homeostasis, mitotic spindle formation and glucose metabolism. While tankyrases are present in most animals, functional differences across species exist. In this work we confirm the widespread distribution of tankyrases throughout the branches of multicellular animal life and identified the single-celled choanoflagellates as earliest origin of tankyrases. We further show that sequence and structural aspects of TNKSs are well conserved even between highly diverged species. We also experimentally characterized an anciently diverged tankyrase homolog from the sponge *Amphimedon queenslandica* and show that the basic functional aspects, such as poly-ADP-ribosylation activity and interaction with the canonical tankyrase binding peptide motif, are conserved. Conversely, the presence of tankyrase binding motifs in orthologs of confirmed interaction partners vary greatly between species, indicating that tankyrases have different sets of interaction partners depending on the animal lineage. Overall, our analysis suggests a remarkable degree of conservation for tankyrases, although their regulatory functions in cells have likely changed considerably throughout evolution.

## Introduction

Tankyrases (TNKSs) are part of the ARTD enzyme family and they catalyze a sequential transfer of ADP-ribose from NAD^+^ to their protein substrates leaving them modified with poly-ADP-ribosyl (PAR) chains (Haikarainen et al., 2014a; Smith et al., 1998). TNKSs regulate a wide variety of pathways and cellular processes, such as telomere maintenance, glucose metabolism, spindle formation, Wnt/β-catenin signalling and Hippo/YAP-signaling (Haikarainen et al., 2014a; Hsiao and Smith, 2008). Humans and other vertebrates possess two highly similar TNKS paralogs: TNKS1 and TNKS2. It is yet unclear if these paralogs have different functions in the cell, however they are at least redundant to some degree as indicated by knockout experiments in mice (Chiang et al., 2008).

Among ARTD members, which are also referred to as PARPs (Lüscher et al., 2021), TNKSs have a unique domain architecture (**Figure 1a**). In addition to their catalytic ADP-ribosyltransferase (ART) domain, TNKSs have five N-terminal ankyrin-repeat (ANK) domains, which are also termed ankyrin repeat clusters (ARCs) and numbered ARC1 to ARC5. Except for ARC3, which may have a structural role (Eisemann et al., 2016; Guettler et al., 2011; Morrone et al., 2012), the ARCs bind interaction partners of TNKSs. These interaction partners bind via a small linear TNKS binding motif (TBM). TBM have a canonical sequence Rxx[ACGP]xGxx, although also less constrained variations such as Rx(4)xGxx and Rx(5)xGxx were reported to bind to TNKSs (DaRosa et al., 2018; Guettler et al., 2011; Huang et al., 2009; Morrone et al., 2012). A sterile alpha motif (SAM) domain is located between the ARC and ART domains and mediates the multimerization of TNKSs (DaRosa et al., 2016; Mariotti et al., 2016; Riccio et al., 2016; Rycker and Price, 2004). In contrast to TNKS2, TNKS1 has an additional and likely disordered N-terminal extension of unknown function which is rich in histidine, proline and serine residues and was accordingly termed an HPS region (Kaminker et al., 2001).

**Figure 1:**
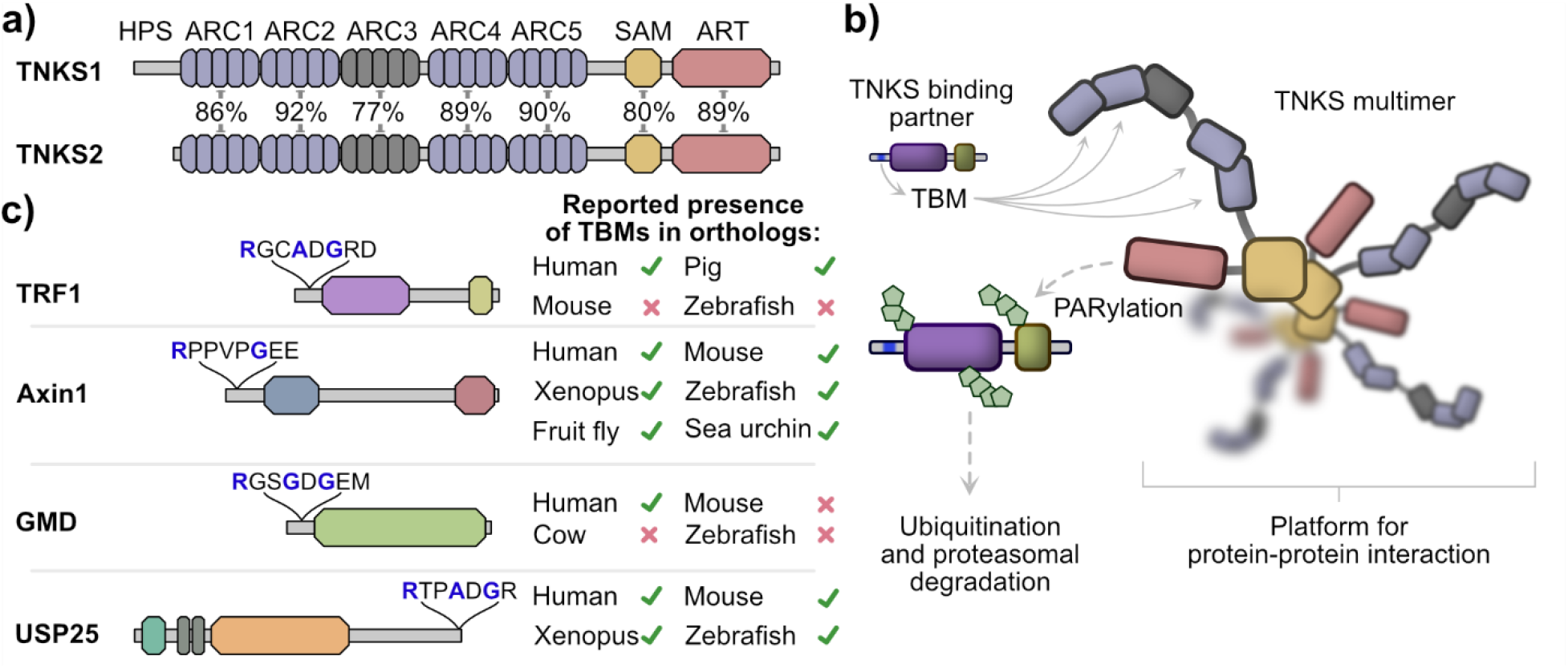
Tankyrase domain organization, function and binding partners. (a) Schematic representation of the domain architecture from human TNKS1 and TNKS2. The percentage sequence identity for domains of TNKS1 and TNKS2 are shown. TNKS1 has a Histidine, Proline and Serine-rich region (HPS) at the N-terminus, which is not present in TNKS2. (b) A model of TNKS function. TNKSs form multimers mediated by the SAM domain and interact with proteins that contain TNKS binding motifs (TBMs), which bind to the ARC1, 2, 4 and 5 domains of TNKSs. Many binding partners are PARylated by TNKSs, which leads to their subsequent ubiquitination and proteasomal degradation. Additionally, TNKSs may also act as scaffolding proteins by providing a platform for protein-protein interactions. (c) Examples of tankyrase binding partners TRF1, Axin1, GMD and USP25. The TBM sequence for each binding partner is shown. Crucial residues for the interaction with TNKS ARC domains are highlighted in blue. Previously reported presence or absence of the TBM in orthologs from other species is shown for TRF1 (Sbodio and Chi, 2002), Axin1 (Huang et al., 2009), GMD (Bisht et al., 2012) and USP25 (Xu et al., 2017).

In the current model, PARylation of proteins binding to TNKSs leads to their ubiquitination and subsequent proteasomal degradation (**Figure 1b**). Additionally, TNKSs may act as scaffolding proteins by providing a platform for protein-protein interactions (Li et al., 2017; Mariotti et al., 2016; Rycker and Price, 2004). Importantly, the regulatory functions that TNKSs exert on different pathways in the cell are a consequence of specific binding partners interacting with TNKSs. Numerous proteins interacting with TNKSs have been reported in humans, although it has become clear these interactions are conserved in varying degrees between species (**Figure 1c**). For example, TNKSs regulate telomer length homeostasis through the interaction with the telomeric protein TRF1 (Cook et al., 2002; Smith and de Lange, 1999). This interaction and thus the function of TNKSs in telomere homeostasis is absent in mice due to a missing TBM in the mouse TRF1 ortholog (Muramatsu et al., 2007; Sbodio and Chi, 2002). In contrast, the function of TNKSs as positive regulators of Wnt signaling through interaction and regulation of Axin levels appears well-conserved across species, and presence of the TBM sequence in Axin orthologs was reported even in invertebrates (Huang et al., 2009). The regulatory function of TNKSs on Wnt signaling was shown in several species such as human (Huang et al., 2009), mouse (Qian et al., 2011), zebrafish (Kuusela et al., 2016) and fruit fly (Feng et al., 2014).

While many interaction partners of TNKSs have been confirmed in humans (Haikarainen et al., 2014a; Hsiao and Smith, 2008; Kim, 2018; Zamudio-Martinez et al., 2021), the presence of TBMs in orthologs of other species was only shown in some cases and to varying degrees in terms of taxonomic coverage (**Figure 1c**). In this work, we examined the distribution of TNKSs across species, the conservation in terms of sequence and structure as well as the conservation of the presence of TBMs in orthologs of 20 human binding partners. We show that TNKSs are widely distributed throughout multicellular animals and we further identify single-celled choanoflagellates as the earliest origin of TNKSs. We demonstrate further that TNKSs display a high degree conservation and show through *in vitro* experiments with an early diverged TNKS ortholog from the demosponge *Amphimedon queenslandica* that the basic molecular functions of TNKSs are also highly conserved. On the other hand, the interaction with different TNKS binding partners appears to be far less conserved, as the presence of TBMs in orthologs vary greatly throughout metazoan evolution.

## Results

### Distribution and origin of tankyrases

Species that are frequently mentioned in this manuscript are shown listed in **Table 1**. We started our analysis by characterizing the distribution of tankyrases throughout the tree of life (**Figure 2a**). Presence of TNKSs was searched within major phylogenetic groups. An early study on the evolutionary history of PARP proteins by Citarelli et al. only found canonical TNKSs present in bilaterians, while Perina et al. later identified TNKSs also in cnidarians and sponges, concluding that TNKSs may be confined to metazoans (Citarelli et al., 2010; Perina et al., 2014). Indeed, we found and confirmed the presence of at least one TNKS homolog in all vertebrates analyzed. While TNKS homologs are present in early diverged bilaterians such as the fruit fly *Drosophila melanogaster* and the mollusk *Octopus bimaculoides*, no TNKS homolog was found in the nematode *Caenorhabditis elegans*. The absence of TNKSs and other PARP homologs in *C. elegans* was reported previously, and the loss of these PARP genes was discussed (Citarelli et al., 2010). It is worth to mention that TNKSs appear present in other nematodes such as *Brugia malayi* and *Trichinella spiralis*.

**Table 1:**
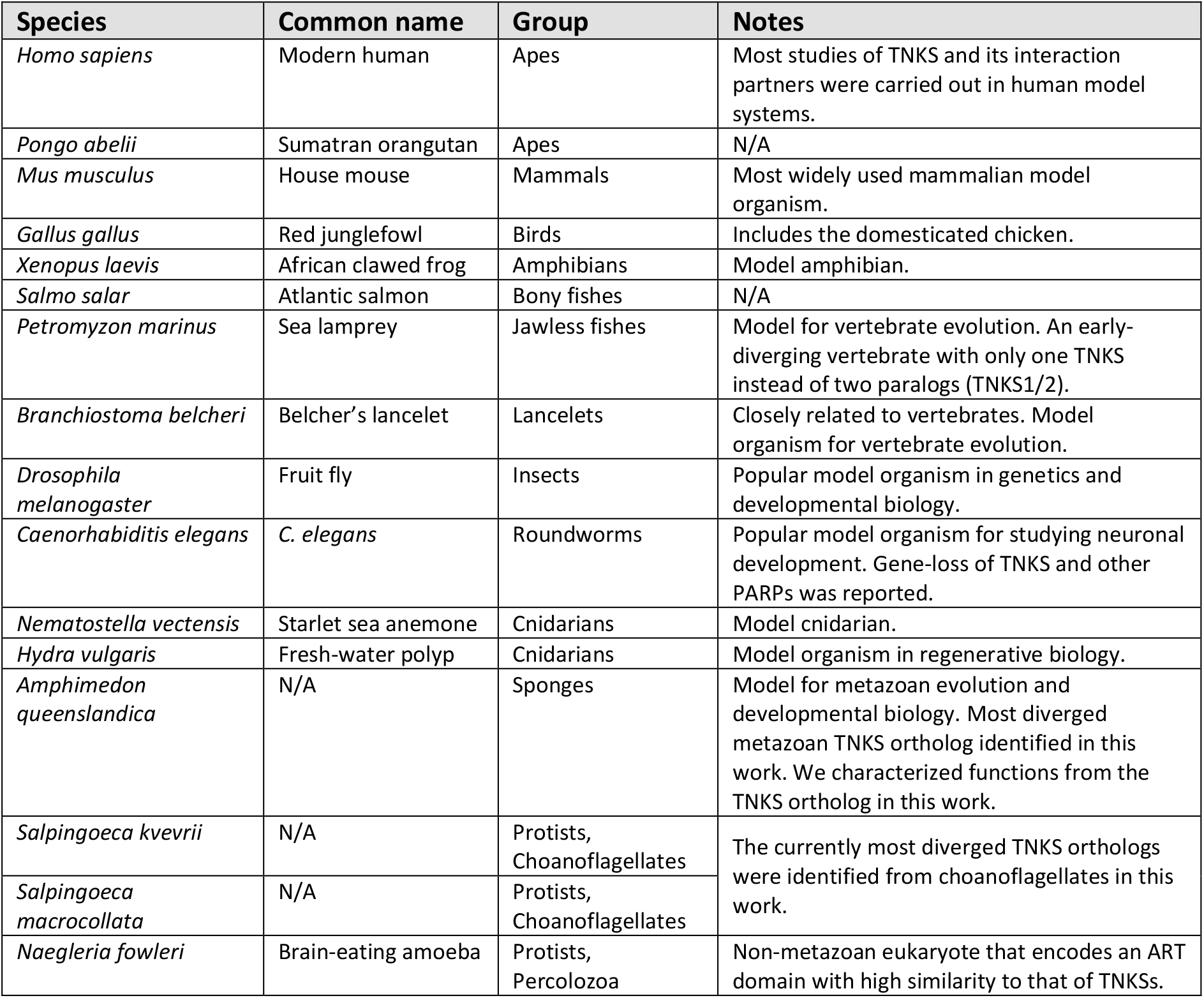
Species frequently referred to in this work.

**Figure 2.**
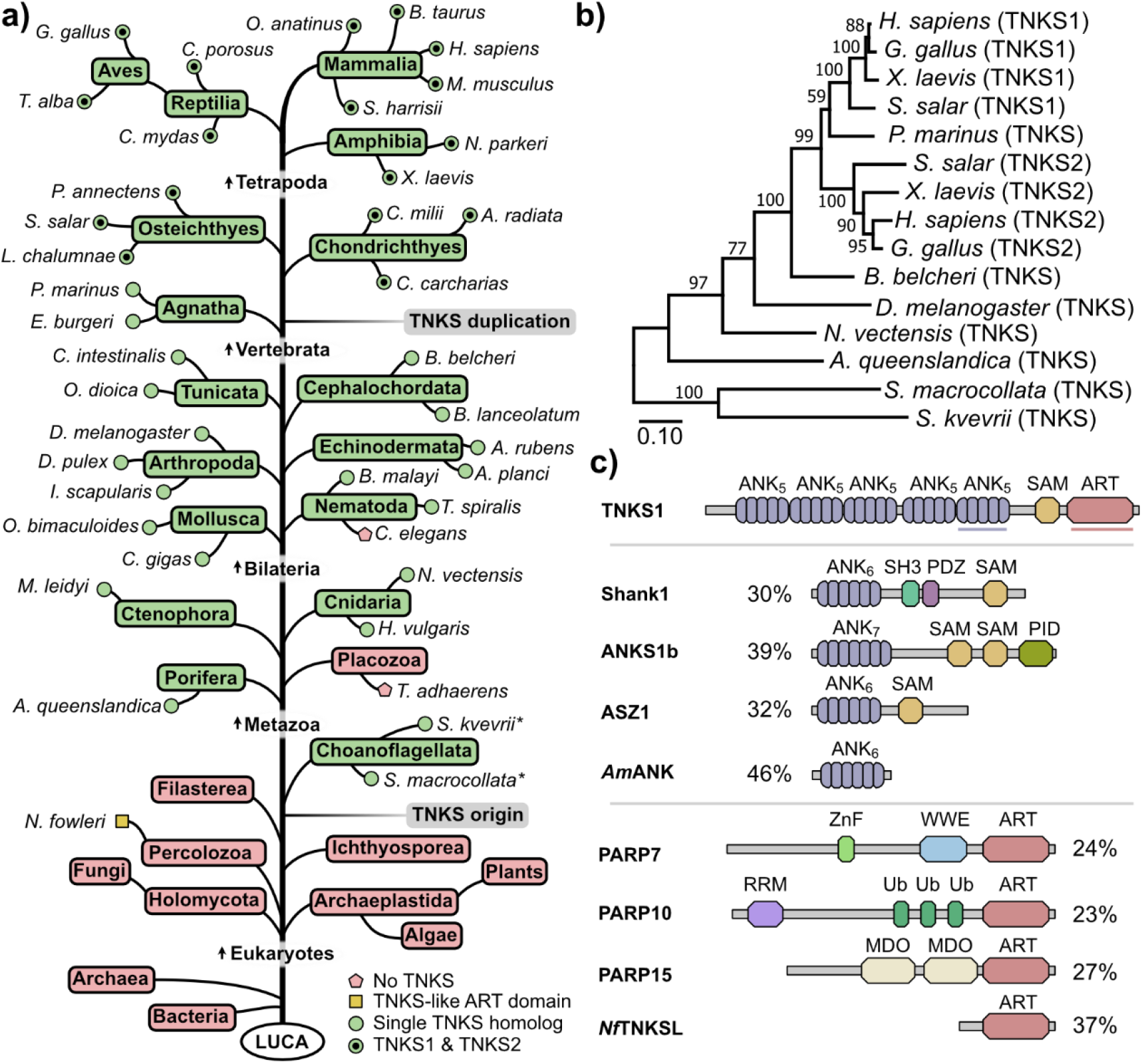
Distribution and possible origin of tankyrases. (a) Presence or absence of TNKSs in phylogenetic groups as determined from searches of various protein and genomic databases. Phylogenetic relationships among groups are schematically shown. Representative species are shown for metazoan and selected groups and the presence of a single TNKS or both TNKS1/2 paralogs is indicated. Information with common names for the phylogenetic groups and species displayed here together with identifiers for the protein sequences can be found in **Table S1**. * = Species in which TNKSs were identified only from TSA data. LUCA = Last universal common ancestor (b) Phylogenetic reconstruction from TNKS sequences using the maximum likelihood method. Percentages for bootstrap support values are shown for each node. Bootstrap tests were done with 1000 bootstrap replicates. The analysis was performed with truncated TNKS sequences starting from the ARC3 domain to match the likely incomplete N-terminal sequence from *S. macrocollata*. In **Figure S1**, the reconstruction was alternatively performed with the neighbor joining method. (c) Schematic representation of proteins with domain architectures related to TNKSs. Sequence identity percentages to the ARC5 or ART domain of human TNKS1 were derived from local sequence alignments. Shank1, ANKS1b and ASZ1 are human proteins that contain both ANK and SAM domains. The ANK protein from *Acidianus manzaensis* (*Am*ANK) shares a high sequence identity with ARC5 of TNKS1. The number of ankyrin repeat units is indicated for each ANK domain. PARP7, PARP10 and PARP15 are human representatives from the clade 3 of the ARTD family, which is most closely related to clade 4 (TNKSs) (Citarelli et al., 2010). The TNKS-like protein from the amoeba *N. fowleri* (*Nf*TNKSL) is described in more detail in **Figure S3**.

Furthermore, our search confirmed that TNKSs are present in early-diverging metazoan species such as the sponge *Amphimedon queenslandica*, the cnidarians *Hydra vulgaris* and *Nematostella vectensis* and the ctenophore *Mnemiopsis leidyi*. However, we could not identify a TNKS homolog in the placozoan *Trichoplax adhaerens*. While the exact taxonomic placement of placozoans is still debated, it is widely accepted that they diverged after the appearance of sponges (Laumer et al., 2018; Srivastava et al., 2008), which indicates a loss of TNKS genes in this lineage similarly to *C. elegans*. In fact, it was suggested that many physiological traits such as muscles, nerves and guts were lost in placozoans in favor of a simpler lifestyle (Laumer et al., 2018; Miller and Ball, 2008), which likely resulted in the loss of many genes.

It has been suggested that the gene duplication event leading to TNKS1 and TNKS2 occurred at some point during the evolution of fishes (Perina et al., 2014). Our analysis shows that the two TNKS paralogs can be identified in representatives of bony fishes (Osteichthyes) and cartilaginous fishes (Chondrichthyes), but not in the jawless fishes *Petromyzon marinus* and *Eptatretus burgeri*, indicating that the duplication event took place during the early evolution of vertebrates.

To interrogate a possible pre-metazoan origin, we searched for the presence of TNKSs in representative subsets of earlier diverging eukaryotes. No TNKSs were identified in the major eukaryotic groups Archaeplastida and Holomycota, which include plants and fungi, respectively. We then searched for the presence of TNKS in groups more closely related to metazoans. While we did not find TNKSs in Filasterea or Ichthyosporea, we identified sequences coding for TNKSs in transcriptome shotgun assembly (TSA) data from the choanoflagellates *Salpingoeca kvevrii* and *Salpingoeca macrocollata*. Choanoflagellates are the proposed sister group of metazoans and are studied as models for pre-metazoan evolution along with their ability to live in communities and differentiate into different cell types (Carr et al., 2008; King et al., 2008). It is worth mentioning that we did not detect the presence of TNKSs in the model choanoflagellates *Monosiga brevicollis* and *Salpingoeca rosetta*, for which the genomes have been sequenced (Fairclough et al., 2013; King et al., 2008). However, these model choanoflagellates are relatively closely related and do not sufficiently represent the genetic diversity found among choanoflagellates (Richter et al., 2018). A phylogenetic tree constructed from the TNKS protein sequences of several metazoan species as well as the choanoflagellates *S. kvevrii* and *S. macrocollata* reproduced the expected phylogenetic relationships (**Figure 2b, Figure S1**). These results suggest an origin of TNKSs in choanoflagellates.

Although several proteins such as Shank1, ANKS1b and ASZ1 are known to comprise combinations of ANK and SAM domains (**Figure 2c**), this domain architecture appears unique to TNKSs among PARPs. We speculated that gene-fusion or exon-shuffling (Nagy and Patthy, 2011) between an ancestral PARP and an ANK-SAM protein could represent obvious mechanisms by which TNKSs may have originated. To see if tangible TNKS precursor proteins could be identified, we searched with sub-domain sequences from human TNKS1 against non-metazoan sequence databases.

Surprisingly, we identified proteins with ANK domains that share 40-50% sequence identity with the ARC5 domain of TNKS1 in diverse phylogenetic groups including prokaryotes. However, the high similarity may be explained by the fact that many residues in ANK-folds are well-conserved irrespective of the domain’s function (Mosavi et al., 2004). Indeed, closer inspection of the hit proteins such as the ANK domain from the archaeon *Acidianus manzaensis* (**Figure 2c**) showed that binding of TBM peptides in a similar mode to the TNKS ARC domains would not be possible (**Figure S2**).

From a similar search using the TNKS1 ART domain as the query, we identified a yet uncharacterized protein from the pathogenic amoeba *Naegleria fowleri* that shares 37% sequence identity with the ART domain of TNKS1 (**Figure 2c**). Further analysis indicates that the TNKS-like ART domain from *N. fowleri* (*Nf*TNKSL) possesses a zinc-binding motif corresponding to the motif in the ART domain of TNKSs (**Figure S3**). It was thought that this motif was unique to TNKSs among PARPs (Lehtiö et al., 2008), and we showed recently that it is important for the structural makeup of the TNKS ART domain (Sowa and Lehtiö, 2022). The available sequence information of *Nf*TNKSL indicates that it does not contain other domains. Considering the similarity, *Nf*TNKSL likely shares a common ancestor with TNKSs and it may resemble an ancestral version of TNKSs before acquisition of ANK and SAM domains.

### Conservation of tankyrase sequence and structure

The results above confirm the vast distribution of TNKSs among metazoan groups and additionally in choanoflagellates. We next aimed to characterize how well-conserved TNKSs are throughout evolution in terms of sequence, structure and function. The following analyses were performed in respect to metazoans with well-characterized genomes; choanoflagellates were omitted from the analyses as their sequence information originated from TSA and may thus be incomplete and less reliable.

First, we compared the sequence identity of human TNKS1/2 and other human ARTD family members with their orthologs from representative metazoan species derived from multiple sequence alignments (**Figure 3**). For further comparison, the ubiquitous and well-conserved proteins cytochrome C (Baba et al., 1981) and seryl-tRNA synthetase (Härtlein and Cusack, 1995) were included as controls. It was reported that some PARPs such as PARP4 and PARP9 have experienced rapid evolution likely due to positive selection in response to host-virus conflicts (Daugherty et al., 2014). Indeed, PARP4, PARP9 and PARP12 share less than 50% sequence identity between the homologs of humans and the fish *S. salar*. In contrast, TNKSs showed the highest degree of conservation of any analyzed family member, displaying approximately 60% sequence identity between human TNKS1/2 and the distant TNKS ortholog found in the demosponge *A. queenslandica*. The degree of conservation appears comparable to that of cytochrome C and seryl-tRNA-synthetase, suggesting that TNKSs are highly conserved members of the ARTD family. Remarkably, the level of conservation for TNKSs within the metazoan lineage appears greater than that for PARP1, which is known to act as key-player in DNA damage response (Pascal, 2018) and its orthologs are widely distributed throughout eukaryotes in protists, animals, fungi and plants (Perina et al., 2014).

**Figure 3:**
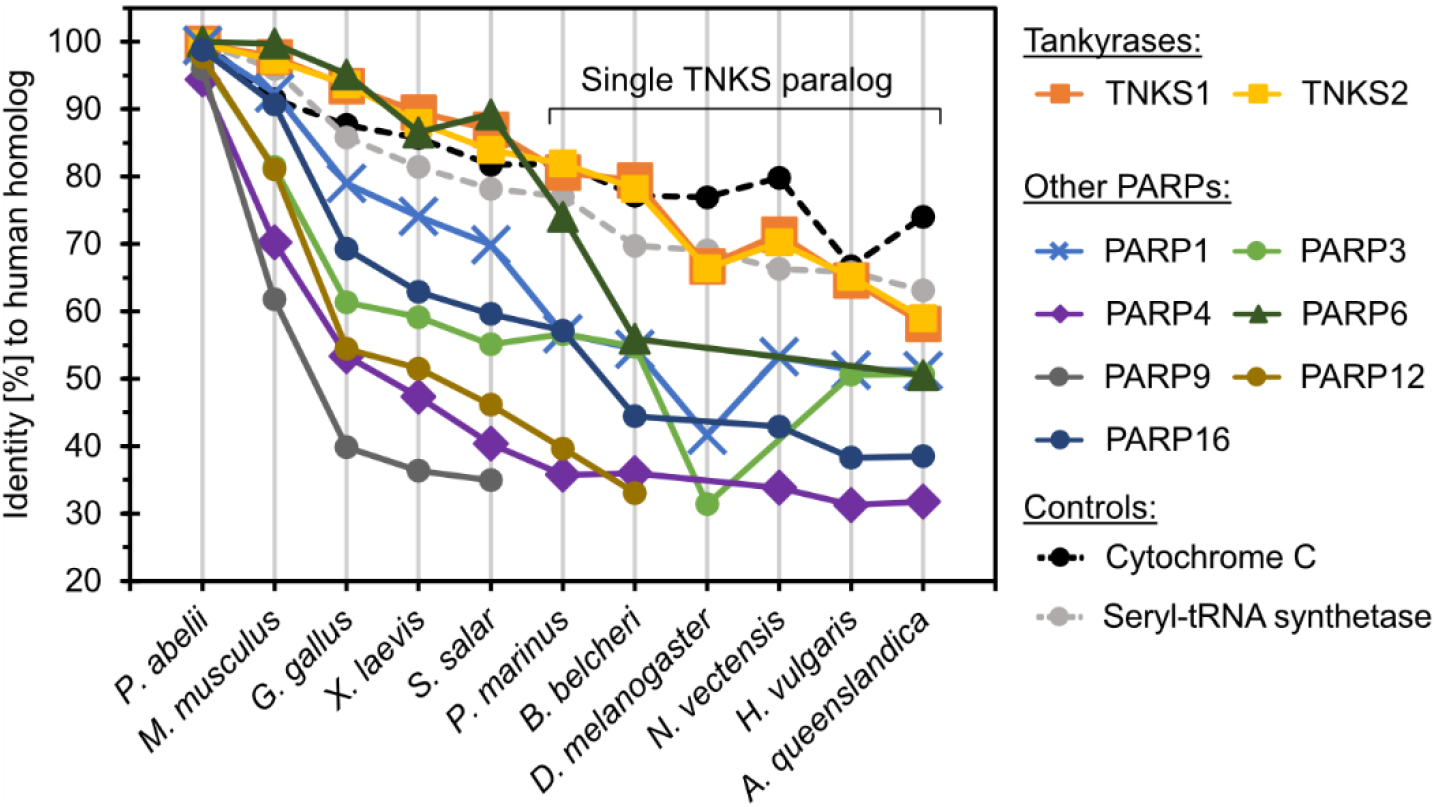
Sequence conservation of tankyrases and representative PARP proteins. For the different proteins analyzed, the graph shows the sequence identity shared by the human ortholog with orthologs from different metazoan species. The identities were determined from multiple sequence alignments of the orthologs from each protein, respectively. In species that only encode a single TNKS paralog, the sequence identity shared with human TNKS1 or TNKS2 was determined against the same TNKS ortholog. The conserved proteins cytochrome C and seryl-tRNA synthase were used for comparison.

To gain insight into the localization of potentially highly or poorly conserved regions in TNKSs, we plotted the number of different residues for each position from an alignment of 113 TNKS sequences from taxonomically diverse metazoans (**Figure 4a**). We additionally used ConSurf analysis, which provides normalized grades for the conservation at each position (Ashkenazy et al., 2016). Because the N-terminal region in TNKS1 is poorly conserved among species, we used human TNKS2 as reference protein to avoid ambiguity due to insufficient alignments in this region.

**Figure 4:**
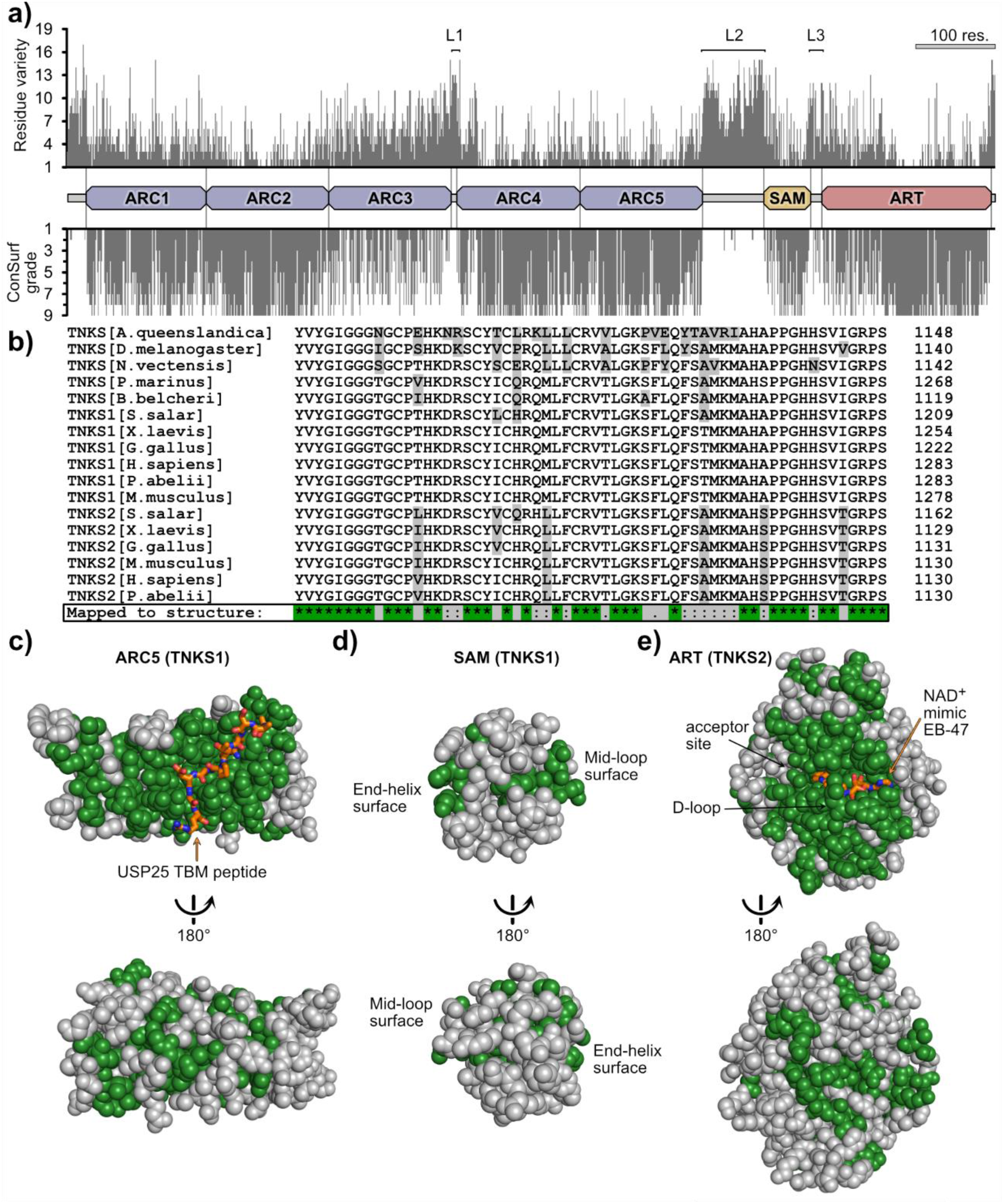
Conservation of tankyrase residues linked to the protein structure. A) Residue-specific conservation plot. The number of different residue identities and ConSurf grade was plotted for each residue from an alignment of 113 TNKS sequences. The ConSurf grade is a normalized measure for the conservation and ranges from 1 (low conservation) to 9 (high conservation). The three likely flexible linker regions are designated L1, L2 and L3. The projected domain boundaries were assigned based on existing structural information of individual domains from human TNKS1 and TNKS2. (b) A representative section of a multiple sequence alignment from the TNKS sequences of 11 metazoan species. This section covers a part of the ART domain. Residues highlighted in gray are different compared to human TNKS1. Only residues identical in every TNKS sequence were mapped to the structures and are shown in green: (c) Structure of the ARC5 domain from human TNKS1 in complex with the TBM peptide from USP25 (orange) (PDB: 5GP7, (Xu et al., 2017)). (d) Structure of the SAM domain from human TNKS1 (PDB: 5KNI, (Riccio et al., 2016)). (e) Structure of the ART domain from human TNKS2 in complex with the NAD^+^-mimicking compound EB-47 (orange) (PDB: 4BJ9, (Haikarainen et al., 2014b)).

This analysis shows strong conservation for the ARC domains 1-2 and 4-5, the SAM domain and the ART domain. The ARC3 domain clearly shows a lower degree of conservation in comparison to other ARCs. This ARC3 was reported not to bind TBMs and likely has a structural role (Eisemann et al., 2016; Guettler et al., 2011; Seimiya et al., 2004). Overall, the lowest degree of conservation is found in the regions linking the domains ARC3 to ARC4, ARC5 to SAM and SAM to ART as well as the N- and C-terminal regions. These regions are likely intrinsically disordered, thus allowing a higher compositional freedom. A study by Eisemann et al. investigated the structural features of the ARC region from TNKS1 and demonstrated that the linking region from ARC3 to ARC4 is indeed flexible, while the regions from ARC1 to ARC2 and ARC2 to ARC3 form rigid interfaces (Eisemann et al., 2016). Although points of flexibility were also observed between ARC4 and ARC5 (Eisemann et al., 2016), our analysis shows a high level of conservation at their interface, possibly indicating that the flexibility does not originate from an intrinsically disordered linker region.

In order to further characterize the sequence conservation of TNKSs in respect to their structures, we mapped strongly conserved residues to TNKS domain structures. From an alignment of the TNKS sequences from 11 metazoan species with well-characterized genomic information (**Figure 4b**), we mapped only identical residues to the domain structures of ARC5, SAM and ART (**Figure 4c-e**).

The ARC domains 1, 2, 4 and 5 show a large, continuous patch of highly conserved residues around the TBM-binding pocket (**Figure 4c, Figure S4**), indicating that the ability to bind the same peptide motif is conserved throughout TNKS evolution. Such a conserved patch is missing for ARC3 (**Figure S4**), which is in agreement with its inability to bind TBM peptides in human TNKSs. Indeed, inspection of ARC3 sequences from early-diverging metazoans confirms the absence of crucial residues required for binding TBMs (**Figure S5**), suggesting that the ARC3 domain did not only recently lose the TBM-binding function.

Sequences from the SAM domains have fewer conserved sites in common (**Figure 4d**), however the residues at its interaction sites appear highly conserved. These interaction sites are termed “mid-loop” and “end-helix” and allow the TNKS SAM domains to multimerize via head-to-tail interactions (DaRosa et al., 2016; Mariotti et al., 2016; Riccio et al., 2016). The ART domain shows a high degree of conservation at and in proximity of the NAD^+^-binding site (**Figure 4e**). The binding mode of NAD^+^ is therefore likely the same across TNKSs from different species. The residues of the donor loop (D-loop) are likewise entirely conserved. The D-loop was shown to adapt different conformations and partially occupies the NAD^+^-binding pocket in crystal structures of apo-TNKSs (Haikarainen et al., 2014a); it was speculated to be involved in the activity regulation of TNKSs (Fan et al., 2018). Overall, these results suggest a remarkable level of conservation in the evolution of TNKSs and imply that the molecular functions across TNKSs are likely conserved.

### Basic molecular characteristics of TNKSs are conserved

The human paralogs TNKS1 and TNKS2 are the most studied TNKS proteins. As shown above, the sequence of TNKSs across all metazoa is highly conserved. While the distant TNKS homolog in drosophila was functionally characterized in *in vivo* studies in terms of cellular function (Cho-Park and Steller, 2013; Feng et al., 2014, 2018; Wang et al., 2016), we wanted to confirm that the molecular functions of distantly related TNKS homologs are conserved using *in vitro* studies with purified proteins. For this, we recombinantly produced TNKS proteins of a basal metazoan, the demosponge *A. queenslandica* (*Aq*TNKS), comprising the ART domain, the ART-SAM domains and the ARC5 domain. Corresponding constructs from human TNKS1 were tested for comparison.

First, we tested the poly-ADP-ribosylation activity of the ART and SAM-ART constructs by auto-modification (**Figure 5a**). The proteins were incubated with biotinylated NAD^+^ and the incorporation of biotin into the PAR chains was detected in Western blot using HRP conjugated streptavidin. A smear corresponding to the PARylation of the proteins was clearly detected for the SAM-ART constructs of *Aq*TNKS and TNKS1, but not for the isolated ART domains. Moreover, to detect PAR chains that originate from PARylation activity during the recombinant expression, we used the PAR-binder ALC1 fused to nanoluciferase (Sowa et al., 2021). Similarly, PAR chains were detected only in the SAM-ART constructs, and incubation with PAR hydrolyzing human PARG showed loss of the smear. These results are in agreement with previous reports showing a significant loss of catalytic activity of human TNKS1/2 in absence of the SAM domain (Fan et al., 2018; Levaot et al., 2011; Mariotti et al., 2016; Riccio et al., 2016; Rycker and Price, 2004). Therefore, the role as poly-ADP-ribosyltransferases and the SAM-dependent activation mechanism are likely conserved among TNKSs. Moreover, size-exclusion experiments with AqTNKS constructs showed formation of multimeric species for SAM-ART protein but not the ART domain, which is in agreement with the reported multimerization function of the SAM domain in human TNKSs (**Figure S6**) (DaRosa et al., 2016; Mariotti et al., 2016; Riccio et al., 2016; Rycker and Price, 2004).

**Figure 5:**
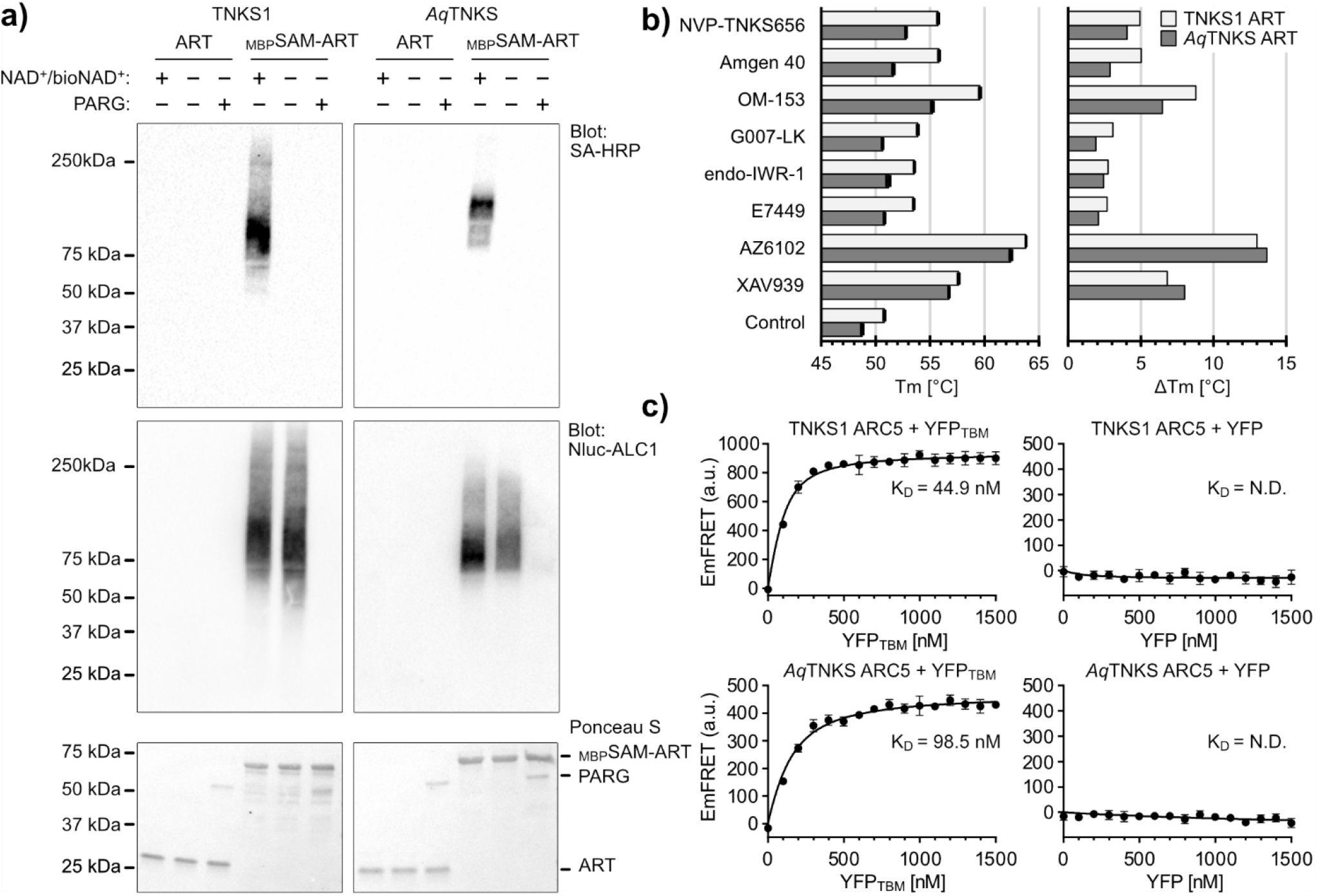
Experimental validation of the tankyrase molecular functions from *Amphimedon queenslandica*. (a) Western blot of ART and SAM-ART constructs from human TNKS1 and *A. queenslandica* TNKS (*Aq*TNKS). SAM-ART constructs have an N-terminal maltose-binding protein (MBP) tag. TNKS constructs (2 μM) were incubated for 1 h at room temperature in presence or absence of a mixture containing biotin-NAD^+^/NAD^+^ (1:10). The total NAD^+^ concentration was 2 μM for TNKS1 and 10 μM for *Aq*TNKS due to a lower activity observed in *Aq*TNKS. PAR with incorporated biotin was detected with streptavidin conjugated to horseradish peroxidase (SA-HRP), while total total PAR modification independent of biotin was detected using nanoluciferase fused to ALC1 (Nluc-ALC1). PARG (0.2 mg/ml) was added to verify removal of PAR chains. (b) Thermal stabilization of TNKS ART domain constructs by selective TNKS inhibitors. Differential scanning fluorimetry (DSF) was performed after mixing TNKS ART domain constructs (5 μM) and TNKS inhibitors (25 μM). A control without inhibitor was included. The absolute melting temperatures (Tm) are shown on the left and data shown are mean ± standard deviation with number of replicates n = 4; the melting temperature relative to the respective controls calculated from the mean values are shown on the right. (c) FRET-based determination of the dissociation constant K_D_ for ARC5 constructs with a TBM peptide. CFP-fused ARC5 constructs (100 nM) from TNKS1 or *Aq*TNKS were mixed with different concentrations of YFP fused to the TBM peptide (REAGDGEE). As control, the CFP-ARC5 constructs were mixed with YFP without the TBM peptide. The fluorescence emissions from FRET (EmFRET) were determined by a method described by Song et al. (Song et al., 2012). Data shown are mean ± standard deviation with number of replicates n = 4.

To further demonstrate the high similarity between the ART domains of *Aq*TNKS and TNKS1, we tested the thermal stabilization with established potent and selective TNKS inhibitors that were recently benchmarked (Brinch et al., 2022). This included adenosine site binders IWR-1, G007-LK, OM-153 and compound 40 (Bregman et al., 2013; Chen et al., 2009; Leenders et al., 2021; Voronkov et al., 2013), nicotinamide-site binders XAV939, E7449, AZ6102 (Huang et al., 2009; Johannes et al., 2015; McGonigle et al., 2015) and the dual-site binder NVP-TNKS656 (Shultz et al., 2013). For all inhibitors, an increased melting temperature for both TNKS1 and *Aq*TNKS was observed (**Figure 5b**), underlining the structural similarity between both ART domains.

Next, we tested the binding of a canonical TBM peptide to the ARC5 domain constructs of *Aq*TNKS and TNKS1. Guettler et al. developed an optimized peptide (sequence: REAGDGEE) that possesses high binding affinity to the ARC domains of human TNKSs (Guettler et al., 2011; Sowa et al., 2020). We expressed the ARC5 domain constructs and the TBM peptide as fusion proteins with the fluorescent proteins CFP and YFP, respectively. To study the interaction of the ARC5 constructs with the TBM peptide, we measured FRET emissions upon mixing of the constructs (**Figure 5c**). Both ARC5 constructs from TNKS1 and *Aq*TNKS interact with the TBM fused to YFP with similar binding affinities, while they showed no interaction with isolated YFP. In combination with the high conservation of the residues lining the ARC5 domain, these results suggest that the ability of TNKSs to bind the same TBM sequences is highly conserved.

### Acquisition of tankyrase binding partners throughout evolution

Experimental data supporting *bona fide* interaction for many of the reported TNKS binding partners in humans exists, although only limited information is available for the interaction of proteins with TNKS orthologs in other metazoan species (Feng et al., 2014). Considering the strong conservation of TNKSs and the ability of *Aq*TNKS to bind the canonical TBM motif RxxϕxGxx, it is reasonable to assume that TNKSs throughout the metazoan lineage can bind this motif. The conservation of the TBM in orthologs of binding partners of human TNKSs has been reported for selected binders, covering often only a very limited taxonomic breadth (Bisht et al., 2012; Huang et al., 2009; Sbodio and Chi, 2002; Xu et al., 2017).

To gain insight into the conservation of TNKS binding partners, we selected a set of 20 experimentally confirmed TNKS interaction partners (**Table 2**) and determined the presence of TBMs in the orthologs of several metazoan species (**Figure 6**). In addition to the canonical TBM Rxx[ACGP]xGxx, we also marked the presence of the less constrained motifs Rx(4)Gxx and Rx(5)Gxx, which were reported for some TNKS interaction partners in humans (DaRosa et al., 2018; Huang et al., 2009; Morrone et al., 2012), although they may require multiple motifs in the target to achieve sufficient avidity. Moreover, we cannot exclude the possibility that despite an apparent lack of TBMs in this analysis, certain orthologs may still interact with TNKSs, for example due to the presence of non-canonical motifs that bind TNKSs.

**Table 2:**
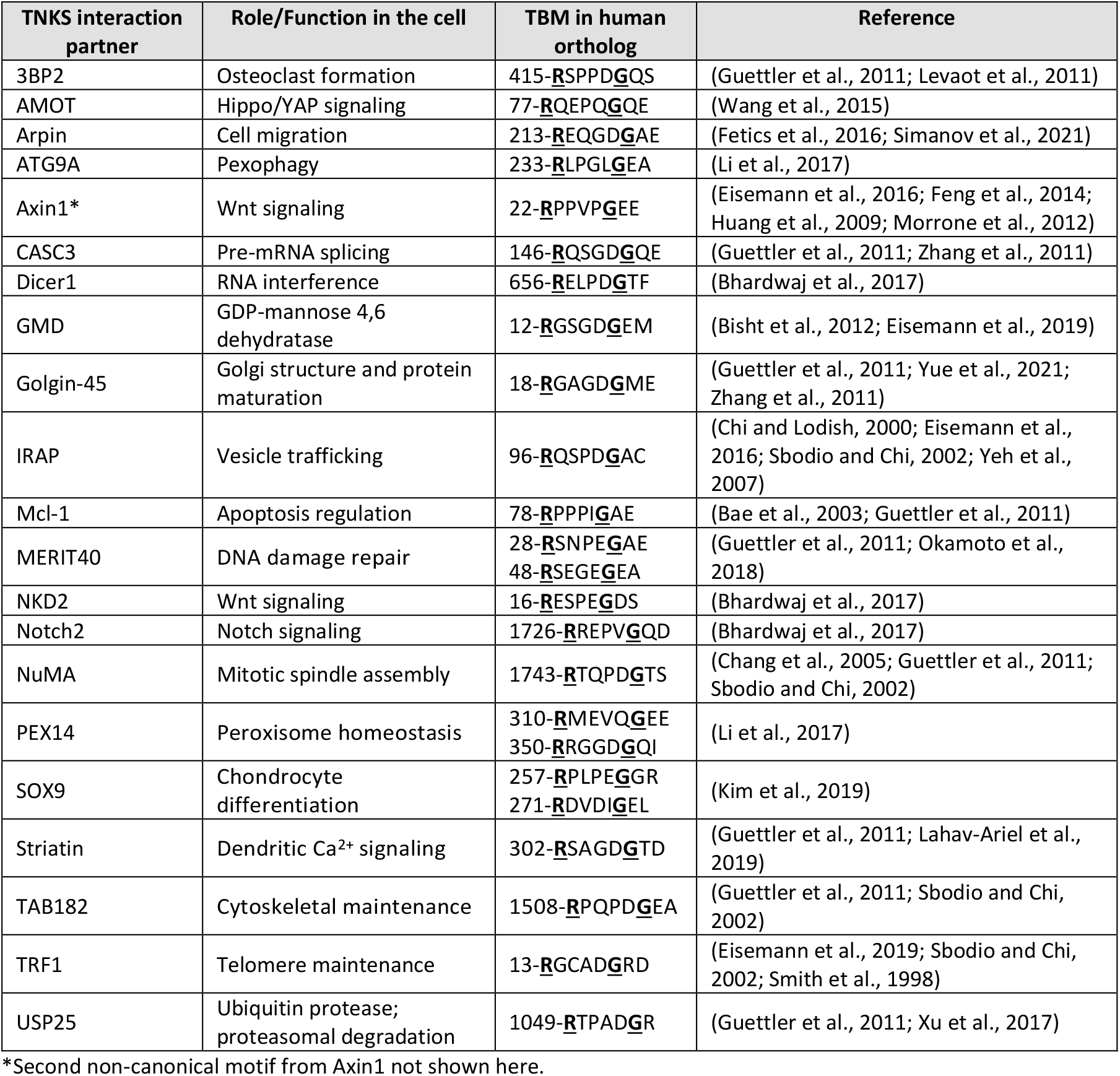
Overview of tankyrase interaction partners used for analysis. The required arginine and glycine in each TBM is shown in bold and underlined.

**Figure 6:**
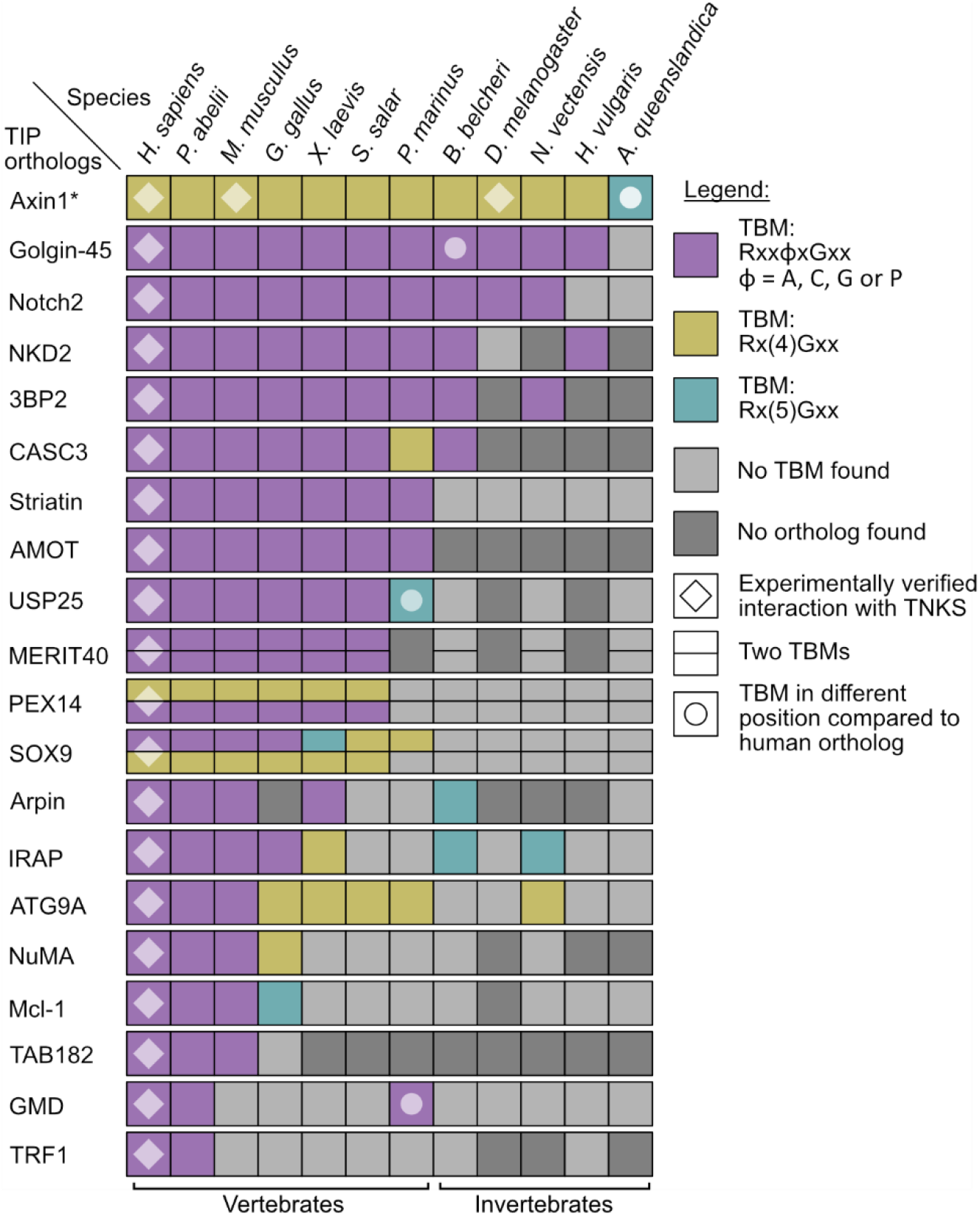
Identification of potential tankyrase binding motifs in orthologs of confirmed interaction partners. Orthologs of 20 TNKS interaction partners (TIPs) were examined for presence or absence of a canonical TBM sequence. Three configurations Rxx[ACGP]xGxx, Rx(4)xGxx and Rx(5)xGxx were considered as TBM. TBMs are only shown if the motif aligned with the TBM in the human ortholog in a multiple sequence alignment, although in some cases also TBMs in nearby positions are shown (see legend). *The non-canonical second motif from Axin1 was not included in this analysis.

The analysis revealed that the degree of conservation of TBMs varies greatly for different binding partners, and some orthologs appeared completely absent in several species. A few of the analyzed orthologs seem to have acquired the TBMs relatively recently, such as GDP-mannose 4,6-dehydratase (GMD) or the telomere length regulator TRF1. Both do not have corresponding TBMs in orthologs of mice or in any of the earlier-diverging metazoans, with the exception of a possible TBM in the GMD ortholog from *P. marinus*. Interestingly, GMD is an otherwise highly conserved enzyme important in basic metabolic pathways and orthologs are present in both prokaryotes and eukaryotes (Somoza et al., 2000). While it was reported that GMD is not a direct target of PARylation by TNKS and moreover appears to inhibit the catalytic activity of TNKSs (Bisht et al., 2012), the exact function of the GMD-TNKS interaction is not well-understood at this point. In contrast, the interaction of TNKSs with Axin1 (Axin) appears highly conserved, as possible TBMs were found in orthologs from all species used for the analysis. Axin is a crucial component in the Wnt signaling pathway and interacts in humans with two TBMs with TNKSs, however we omitted the non-canonical second motif from the analysis as the binding rules for it are not clear (Morrone et al., 2012). Earlier work by Feng et al. also confirmed interaction of TNKS and Axin in *D. melanogaster* (Feng et al., 2014). TBMs in Golgin-45 were identified in orthologs from many of the analyzed species with exception of *A. queenslandica*. Golgin-45 is a Golgi-associated protein required for Golgi structure and protein transport. Zhang et al. identified Golgin-45 as a target for PARylation by TNKSs (Zhang et al., 2011). More recently, it was also shown to be responsible for TNKS1 localization to the Golgi apparatus during interphase (Yue et al., 2021).

The results obtained here lead to the overall picture that TNKSs themselves are highly conserved, while the interaction with binding partners based on the presence of TBMs seems to be far more variable throughout evolution.

## Discussion

The main points derived from this work are the following: First, we showed that TNKSs are distributed widely throughout metazoans, though they appear to have a pre-metazoan origin as demonstrated by the presence in some choanoflagellates. Second, TNKSs appear remarkably conserved in terms of sequence, structure and molecular function. Last, the conservation of TBMs in TNKS binding partners and their orthologs varies greatly, indicating a dynamic gain and possibly a loss of TBMs throughout evolution.

### Origin and distribution

The evolutionary transition from a single-celled urchoanozoan (ancestor of choanoflagellates and metazoans) to metazoans occurred together with the innovation and loss of gene families (Paps and Holland, 2018; Richter et al., 2018). Proteins that originated during this transition and that were retained in the metazoan lineage are of interest for investigations about the evolutionary origin of metazoans, as these proteins may have been important factors for allowing the development of multicellular animals.

TNKSs seem to appear only sparsely in choanoflagellates but are vastly distributed throughout metazoans, which implies that TNKSs are important for multicellular development. However, the presence of TNKSs seems not to be obligatory for metazoan life, as evidenced by the reported loss of TNKS in *C. elegans* (Citarelli et al., 2010; Perina et al., 2014) and possibly in placozoans as shown in this work.

As part of the analysis we identified a yet uncharacterized ARTD family member in the pathogenic amoeba *N. fowleri*, here termed *Nf*TNKSL. Based on the sequence information available, *Nf*TNKSL contains only a single ART domain, which shows more similarity to the ART domain of TNKSs than that of any other ARTD member. It is tempting to speculate that *Nf*TNKSL may resemble an ancestral TNKS precursor, which later obtained ANK and SAM domains through domain-shuffling, a common mechanism for the innovation of novel protein functions in the stem animal lineage (Ekman et al., 2007; López-Escardó et al., 2019; Tordai et al., 2005).

### Conservation of tankyrases

We have shown that the sequences of tankyrases are highly conserved throughout metazoans, displaying similar levels of conservation to fundamentally important proteins such as cytochrome C. As was expected, the degree of conservation is not equal along the TNKS sequence but corresponds to the structure of the protein. For example, regions thought to be flexible linkers appear less conserved, although they are likely important for the function of TNKSs (Eisemann et al., 2016). Mapping of conserved regions from TNKSs to the structures showed a very high degree of conservation particularly around the TBM binding site of ARCs (exemplified for ARC5) and the NAD^+^binding site of the ART domain.

Functional studies comparing human TNKS1 with *Aq*TNKS from *A. queenslandica*, a sponge that diverged from other metazoans over 600 million years ago (Srivastava et al., 2010), confirmed that the basic molecular functions are conserved. We demonstrated that *Aq*TNKS is a poly-ADP-ribosyltransferase, the presence of the SAM domain strongly increased activity, and that the ARC5 domain recognizes and binds a canonical TBM with similar affinity compared to human TNKS1. The ability of *Aq*TNKS to bind structurally diverse TNKS inhibitors that display high selectivity and nanomolar binding affinity against human TNKSs further highlights the high degree of conservation.

### Acquisition of tankyrase binding partners

From an analysis of the presence of TBMs in orthologs of human TNKS binding partners, the number of orthologs with TBMs decreased according to the species divergence from humans. Our analysis indicates that new TBMs formed throughout different animal lineages. This may leave the impression of a trend favoring accumulation of TNKS binding partners as the complexity of animal species increases, however it is important to clarify that a loss of TBMs may have occurred as well. The fact that the vast majority of TNKS binding partners were only experimentally confirmed in humans limits our analysis to the emergence of TBMs in orthologs of these binding partners; a comprehensive analysis for the loss in TBMs would require multiple experimentally verified TNKS binding partners in early-diverging metazoan lineages. From the current results, we cannot exclude the possibility that the number of TNKS interaction partners for example in early-diverging groups like sponges or arthropods may be similar to the number found in humans.

Hub proteins binding small linear motifs (SLiMs) were recently suggested to be grouped under the term of “linear motif-binding hubs” (LMB-hubs) (Jespersen and Barbar, 2020). With TBMs being SLiMs, and considering the functions of TNKSs, they certainly can be classified as LMB-hubs. Evolutionary analyses previously performed on other LMB-hub proteins established striking parallels also observed here: While the hub proteins are generally highly conserved, the presence of interacting SLiMs is often poorly retained in the orthologs of binding partners (Davey et al., 2012; Van Roey et al., 2014). This may be the case because SLiMs can rapidly evolve *de novo* or similarly be lost again through small changes in amino acid sequences (Davey et al., 2015). SLiMs are located on intrinsically disordered protein regions such as the protein termini, which are often far less mutationally constrained than structured regions – allowing rapid changes in sequence composition (van der Lee et al., 2014). Furthermore, SLiM-Hub interactions may allow a level of fine-adjustment for the regulation of binding partners through change of the SLiM sequence, resulting in changes in binding affinity to the hub protein (Davey et al., 2015).

Indeed, it has been reported that differences in the sequence of TBMs result in varied binding affinities to TNKSs (Eisemann et al., 2019; Guettler et al., 2011). A study by Eisemann et al. recently demonstrated that mutations in the TBM of TRF1 reduced the binding affinity and also resulted in lowered PARylation of TRF1 by TNKS1 *in vitro*. Thus, the TNKS-TBM interaction may allow freedom throughout evolution for TNKS binding partners to adapt their TBMs and thus adjust the degree by which they are targeted by TNKS-mediated PARylation. This adjustment of binding may not only happen through changes in the TBM sequences but could also take place through evolution of additional TBMs in the TNKS binding partners or *via* their dimerization, which creates a complex relationship in terms of multivalent binding to the multimeric TNKS scaffolds.

### Cellular and physiological functions of tankyrases

The molecular functions encompassed by TNKSs appear simple: The binding of interaction partners *via* its ARC domains, the PARylation of said partners and formation of multimeric complexes by the SAM domain leading to increased catalytic activity and further providing a binding platform for protein-protein-interactions. In contrast, the cellular or physiological functions of TNKSs and the biological outcomes thereof can be attributed to the diverse assortment of binding partners from many different pathways that are regulated by TNKSs.

While the molecular functions were assessed to be highly conserved, the cellular functions of TNKSs may vary greatly depending on the species as different sets of TNKS binding partners may be regulated by TNKSs. An example of this is TRF1, a nuclear protein repressing telomere elongation which interacts through a TBM with TNKSs in humans. While it was found that TNKSs have an important role in telomere maintenance through regulation of TRF1 in humans (Cook et al., 2002; Dynek and Smith, 2004; Ha et al., 2012; Rippmann et al., 2002; Smith and de Lange, 2000; Smith et al., 1998), the TBM is absent from the TRF1 ortholog in mice, where TNKSs were consistently shown not to participate in the regulation of telomere length (Donigian and de Lange, 2007; Hsiao et al., 2006; Muramatsu et al., 2007). Although the attenuation of telomere elongation through inhibition of TNKSs could be a potential avenue for cancer therapeutics (Seimiya et al., 2005), mouse models would not directly replicate the potential efficacy of such drugs. On the other hand, inhibition of TNKS was shown to reduce Wnt-signaling activity through blocking the PARylation of Axin (Huang et al., 2009), which has a highly conserved TBM across species. Treatment with TNKS inhibitors leads to reduced proliferation of cancer cells in Wnt-dependent tumors (Thorvaldsen, 2017) and mice are frequently used as model organisms in preclinical studies. For the evaluation of TNKS inhibitors as therapeutic drugs, it may need to be considered that the cellular functions TNKSs can differ in humans and model organisms, which may limit for example how well the perceived efficacy or toxicity translates to humans.

## Acknowledgements

Biocenter Oulu Structural Biology core facility, member of Biocenter Finland, Instruct-ERIC Centre Finland and FINStruct, as well as of “Proteomics and Protein Analysis” and Sequencing core facilities are gratefully acknowledged. We also acknowledge the support of the Biocenter Finland Bioinformatics platform node at Åbo Akademi University.

## Supplementary Data

**Supplementary Information.** Contains supplementary Table S1 and supplementary Figures S1-6.

## Methods

### Sequence acquisition

Canonical sequences from human orthologs were obtained from UniProt. Sequences from non-human orthologs were obtained from NCBI non-redundant (NR) database using sequences of human orthologs as a query by BLAST (http://blast.ncbi.nlm.nih.gov/Blast.cgi) (Altschul et al., 1990). If no ortholog could be identified in protein databases, the tBLASTn implementation against the Genebank database (TSA and WGS) was used. Sequences of TNKSs from *E. burgeri* and *M. leidyi* were acquired from the Ensembl database (https://metazoa.ensembl.org/). Possible functional annotations for the TNKS-like protein from *N. fowleri* were examined in the Amoeba database (https://amoebadb.org/) under the gene identifier NF0130940.

### Sequence alignments and construction of phylogenetic trees

Multiple sequence alignments were done with Clustal Omega (Sievers et al., 2011) or the Clustal W implementation in the MEGA11 software (Tamura et al., 2021). Global pairwise sequence alignments were done with EMBOSS Needle and local pairwise alignments were done with EMBOSS Matcher. The construction of phylogenetic trees was done with the maximum likelihood and neighbor joining methods in MEGA11 (Tamura et al., 2021) using the Jones-Taylor-Thornton (JTT) substitution model (Jones et al., 1992) and a bootstrap analysis with 1000 replicates was performed for each tree.

### Structure visualization and prediction

Models of experimentally derived protein structures were obtained from the Protein Data Bank (PDB) (Berman, 2000). The structure models were visualized in the PyMOL Molecular Graphics System (PyMOL, version 1.8.4.0). Prediction of protein structures from *A. manzaensis* and *N. fowleri* were done with RoseTTAFold (Baek et al., 2021) using only the protein sequences as input without additional constraints.

### Sequence-specific conservation analysis

Sequences of TNKSs originating from a broad taxonomic range were sampled and obtained as described above. Sequences annotated as low-quality proteins were removed and the remaining sequences were inspected for obvious indications of assembly errors. A multiple sequence alignment from 113 TNKS sequences was used as input for residue specific conservation analysis by ConSurf (Ashkenazy et al., 2016). JTT was determined to be the best evolutionary model for the data provided and was used by ConSurf to estimate the evolutionary relationships. ConSurf conservation scores were calculated based on the Bayesian method.

### Identification of potential TBMs in orthologs of TNKS binders

Sequences for the orthologs of the human TNKS interaction partners were obtained as described above. Multiple sequence alignments were performed for each set of orthologs, and the presence of potential TBMs in in respect to the human ortholog was examined for each set. The different configurations Rxx[PGAC]xGxx, Rx(4)xGxx and Rx(5)xGxx were accepted as possible TBMs. Generally, only those TBMs that aligned with the position of the TBM in the human ortholog were considered potential TBMs; although not aligned, some possible TBMs were annotated due to presence in a likely disordered region in proximity to the TBM in human orthologs.

### Molecular cloning

The cloning of human TNKS1 constructs was done as previously described for CFP-ARC5 (Sowa et al., 2020) or SAM-ART and ART domains (Sowa and Lehtiö, 2022). Cloning of the YFP-REAGDGEE construct was done as previously described (Sowa et al., 2020). The *E. coli* codon-optimized constructs for *A. queenslandica* TNKS (*Aq*TNKS: XP_019848937.1) were obtained as gblock from IDT. The construct ARC5 (*Aq*TNKS Glu658-Met811) was cloned into the pNIC-CFP expression vector (Addgene #173074). The *Aq*TNKS constructs SAM-ART (Pro889-Thr1181) and ART (Thr968-Thr1181) were cloned into the pNIC-MBP vector, which was prepared as previously described (Sowa et al., 2020). Cloning was done using the SLIC method (Jeong et al., 2012). Briefly, the vectors were linearized and mixed with insert and T4 DNA polymerase. The reaction mixture was used to transform NEB 5α chemicompetent *E. coli* cells (New England BioLabs). Colonies were grown at 37°C on LB agar plates containing 5% sucrose and 50 μg/ml kanamycin; genes in the vector encoding kanamycin-resistance and SacB enzyme (Hynes et al., 1989; Reyrat et al., 1998) served as selection markers for successful transformation and vector linearization, respectively. All constructs were verified by sequencing of the insert regions.

### Protein expression

Chemicompetent *E. coli* BL21(DE3) cells were transformed with the expression vectors. For each construct, 500 ml Terrific Broth (TB) autoinduction media including trace elements (Formedium, Hunstanton, Norfolk, England) were supplemented with 8 g/l glycerol and 50 μg/ml kanamycin and inoculated with overnight preculture. The flasks were incubated at 37 °C (shaking) until an OD600 of 1 was reached. The incubation continued overnight at 18 °C (shaking). At the end of the incubation period, the cells were harvested by centrifugation at 5,000 xg, 4°C for 30 minutes. The pellets were resuspended in lysis buffer (50 mM HEPES pH 7.5, 500 mM NaCl, 0.5 mM TCEP, 10 mM imidazole, 10% glycerol v/v) and used immediately for purification or frozen at −20°C.

### Protein purification

Purification of human TNKS1 ART and TNKS1 SAM-ART (Sowa and Lehtiö, 2022) or CFP-ARC5 (TNKS1) and YFP-REAGDGEE (Sowa et al., 2020) was done as previously described. Frozen cells were thawed first. The cells were supplemented with 0.1 mM Pefablock SC (Roche) and 20 μg/ml DNase I (Roche) and lysed by sonication. The lysate was centrifuged (30,000×g, 4°C, 30 min) and filtered (0.45 μm). As initial purification step, all the proteins were purified by immobilized metal affinity chromatography (IMAC) with a 5 ml HiTrap HP column equilibrated with lysis buffer and charged with Ni^2+^. Lysis buffer with higher concentrations of imidazole was used for washing steps (25 mM) and elution (300 mM).

The IMAC purification of the *Aq*TNKS ART domain (MBP-tagged) was followed by buffer exchange with size-exclusion chromatography (SEC) buffer (20 mM HEPES pH 7.5, 350 mM NaCl, 0.5 mM TCEP, 10% v/v glycerol) using an Amicon Ultra-15 centrifugal filter (30 kDa MWCO). To remove MBP, the sample was incubated with TEV protease (1:30 molar ratio) for 48h at 4°C and was then run over a 5 ml MBPTrap HP column equilibrated with SEC buffer. SEC was performed as last step of the purification using S75 16/600 size-exclusion chromatography column and SEC buffer.

For *Aq*TNKS SAM-ART (MBP-tagged), the IMAC step was followed by MBP affinity chromatography using a 5 ml MBPTrap HP column equilibrated with SEC buffer. After loading, the column was washed with 5-10 column volumes of SEC buffer and eluted with SEC buffer containing 10 mM maltose.

For the *Aq*TNKS ARC5 domain (CFP-tagged), the elution from IMAC was purified by SEC as above. To remove contamination by *E. coli* chaperone proteins, a successive IMAC purification was performed and 5mM MgCl and 5 mM ATP disodium salt were included in the washing step (Rial and Ceccarelli, 2002). Finally, a second purification by SEC was performed as above. At the end of each purification, all the proteins underwent concentration, they were aliquoted, flash frozen in liquid nitrogen and stored at −70 °C.

### Activity analysis by Western blot

The constructs of TNKS1 or *Aq*TNKS were prepared in absence or presence of a mixture of biotinylated-NAD^+^/NAD^+^ (1:10). Additionally, a control with PARG (0.2 mg/ml) was prepared. Final total NAD^+^-concentrations were 2 μM and 10 μM for TNKS1 and *Aq*TNKS, respectively. The reactions were incubated for 1 hour at room temperature. At the end of the incubation period, 10 μL of each reaction were run on SDS-PAGE (Mini-Protean TGX 4-20% gradient gel, BioRad). The transfer of the proteins to a nitrocellulose membrane (Trans-Blot Turbo Mini 0.2 μm Nitrocellulose Transfer Pack, BioRad) was performed using a Trans-Blot semi-dry system (BioRad). Afterwards, the membrane was washed with TBS-T and subsequently stained with Ponceau S solution. The membranes were imaged (ChemiDoc Imaging System, BioRad) and the staining was removed by washing the membrane in TBS-T. The membranes were blocked using 1x TBS with 1% casein (BioRad) for 30 minutes. The membranes were incubated with streptavidin-HRP (PerkinElmer) diluted 1:10,000 in blocking buffer for 30 minutes. The membranes were washed with TBS-T buffer and HRP detection was done using ECL solution (Advansta). Following the imaging step, the membranes were washed in TBS-T containing 5%(w/v) skimmed milk powder for 15 minutes. The membranes were transferred to blocking solution including 0.1 μg/ml nanoluciferase-ALC1 (Sowa et al., 2021) for 30 minutes. Finally, the membrane was washed in TBS-T and detection of nanoluciferase on the membrane was done with 1:500 NanoGlo (Promega) diluted in PBS buffer.

### Analytical size-exclusion chromatography

In separate independent runs, 0.5 mg of each protein sample (AqTNKS SAM-ART, AqTNKS ART, TNKS1 SAM-CAT and MBP) was loaded to a Superdex 200 Increase 10/300 GL column. The column was equilibrated in SEC buffer (20 mM HEPES pH 7.5, 350 mM NaCl, 0.5 mM TCEP, 10% glycerol) and samples were run at a flow rate of 0.5 ml/min at 4°C. The absorbance at 280 nm was recorded. In each run, 1 ml fractions were collected. Fractions corresponding to the different peaks observed for the SAM-ART constructs were analyzed by SDS-PAGE.

### Differential scanning fluorimetry

Samples of ART constructs (5 μM) from TNKS1 or *Aq*TNKS were prepared in absence or presence of TNKS inhibitors (25 μM) IWR-1, G007-LK, OM-153, compound 40, XAV939, E7449 or NVP-TNKS656 (Brinch et al., 2022) and 5x SYPRO orange dye (ThermoFisher Scientific). Each condition was prepared in 4 replicates with a final volume of 20 μL. The sample buffer was 10 mM Bis-Tris-Propane (pH 7), 3% PEG 20,000 (w/v), 0.01% Triton X-100 (v/v), 0.5 mM TCEP. The samples were transferred into a 96-well transparent qPCR plate and the measurements were performed using a BioRad C1000 CFX96 thermal cycler. The temperature was increased from 20°C to 95°C (1°C/min) and data points for melting curves were recorded in 1-minute intervals. The data analysis was performed with GraphPad Prism 9 using a nonlinear regression analysis (Boltzmann sigmoid equation) of normalized data.

### Determination of the dissociation constants by FRET

The experiment was performed as previously described (Sowa et al., 2020). Briefly, CFP-fused ARC5 domains (100 nM) from TNKS1 or *Aq*TNKS were mixed with YFP-REAGDGEE (0-1500 μM). The FRET emission for each condition was determined. The calculation of dissociation constants was done using a method described by Song et al. (Song et al., 2012).

