## Supplementary information for "An evolutionary perspective on the origin, conservation and binding partner acquisition of tankyrases"

### Supplementary material content

**Table S1: Protein sequence identifiers from the analysis of tankyrase distribution and origin.**

**Figure S1: Phylogenetic analysis of tankyrases.**

**Figure S2: Predicted structure of the TNKS-like ankyrin-repeat domain from the archaeon *Acidianus manzaensis*.**

**Figure S3: Predicted structure of the TNKS-like ART domain from the amoeba *Naegleria fowleri*.**

**Figure S4: Conservation of TNKS ARC domains mapped to the structures.**

**Figure S5: The ARC3 domains in both human and early-diverging tankyrases are unlikely to bind TBM peptides.**

**Figure S6: Size-exclusion chromatography of TNKS1 and *Aq*TNKS constructs.**

**Table S1: Protein sequence identifiers from the analysis of tankyrase distribution and origin.** Accession identifiers for the NCBI protein database are shown for each species. ANK = Ankyrin-repeat domain; ART = ADP-ribosyltransferase domain. *Data from transcriptome shotgun assembly; identifier refers to the NCBI nucleotide databank. **Identifier refers to the transcript in the EnsemblGenomes database.

| Phylogenetic group | Species | Common name | Accession identifier |
|---|---|---|---|
| Archaea | *Acidianus manzaensis* | N/A | ANK: WP_148690368.1 |
| Percolozoa | *Naegleria fowleri* | Brain-eating amoeba | TNKS-like ART: XP_044559609.1 |
| Choanoflagellata | *Salpingoeca kvevrii* | N/A | TNKS*: GGOX01000296.1 |
| | *Salpingoeca macrocollata* | N/A | TNKS*: GGOT01036632.1 |
| Porifera (Sponges) | *Amphimedon queenslandica* | N/A | TNKS: XP_019848937.1 |
| Placozoa | *Trichoplax adhaerens* | N/A | No TNKS ortholog found. |
| Ctenophora (Comb jellies) | *Mnemiopsis leidyi* | Sea walnut | TNKS**: ML097525a-RA |
| Cnidaria (Cnidarians) | *Nematostella vectensis* | N/A | TNKS: XP_032220531.1 |
| | *Hydra vulgaris* | N/A | TNKS: XP_047128184.1 |
| Mollusca (Molluscs) | *Octopus bimaculoides* | California two-spot octopus | TNKS: XP_014782906.1 |
| | *Crassostrea gigas* | Pacific oyster | TNKS: XP_019923312.2 |
| Nematoda (Roundworms) | *Brugia malayi* | N/A | TNKS: CDP94254.1 |
| | *Trichinella spiralis* | Pork worm | TNKS: KRY42746.1 |
| | *Caenorhabditis elegans* | N/A | No TNKS ortholog found. |
| Arthropoda (Arthropods) | *Drosophila melanogaster* | Fruit fly | TNKS: NP_651410.1 |
| | *Daphnia pulex* | Water flea | TNKS: XP_046457952.1 |
| | *Ixodes scapularis* | Deer tick | TNKS: XP_029842287.2 |
| Echinodermata | *Asterias rubens* | Common starfish | TNKS: XP_033632864.1 |
| | *Acanthaster planci* | Crown-of-thorns starfish | TNKS: XP_022094330.1 |
| Cephalochordata (Lancelets) | *Branchiostoma belcheri* | Belcher's lancelet | TNKS: XP_019641281.1 |
| | *Branchiostoma lanceolatum* | European lancelet | TNKS: CAH1257770.1 |
| Tunicata (Tunicates) | *Ciona intestinalis* | Vase tunicate | TNKS: XP_002121662.3 |
| | *Oikopleura dioica* | N/A | TNKS: CBY09100.1 |
| Agnatha (Jawless fish) | *Petromyzon marinus* | Sea lamprey | TNKS: XP_032806710.1 |
| | *Eptatretus burgeri* | Inshore hagfish | TNKS**: ENSEBUT00000023950.1 |
| Chondrichthyes (Cartilaginous fish) | *Callorhinchus milii* | Australian ghostshark | TNKS1: XP_042200715.1<br>TNKS2: XP_007894887.1 |
| | *Amblyraja radiata* | Thorny skate | TNKS1: XP_032873799.1<br>TNKS2: XP_032889463.1 |
| | *Carcharodon carcharias* | Great white shark | TNKS1: XP_041041558.1<br>TNKS2: XP_041066021.1 |
| Osteichthyes (Bony fish) | *Salmo salar* | Atlantic salmon | TNKS1: XP_014017013.1<br>TNKS2: XP_014034742.2 |
| | *Protopterus annectens* | West African lungfish | TNKS1: XP_043916872.1<br>TNKS2: XP_043912589.1 |
| | *Latimeria chalumnae* | Coelacanth | TNKS1: XP_006007641.1<br>TNKS2: XP_006006371.1 |
| Amphibia (Amphibians) | *Xenopus laevis* | African clawed frog | TNKS1: XP_018099068.1<br>TNKS2: XP_018082988.1 |
| | *Nanorana parkeri* | High Himalaya Frog | TNKS1: XP_018421084.1<br>TNKS2: XP_018427335.1 |
| Reptilia (Reptiles) | *Crocodylus porosus* | Saltwater crocodile | TNKS1: XP_019389515.1<br>TNKS2: XP_019411065.1 |
| | *Chelonia mydas* | Green sea turtle | TNKS1: XP_037752450.1<br>TNKS2: XP_007059469.2 |
| Aves (Birds) | *Gallus gallus* | Red junglefowl | TNKS1: NP_989671.2<br>TNKS2: NP_989672.2 |
| | *Tyto alba* | Barn owl | TNKS1: XP_042664360.1<br>TNKS2: XP_032840041.2 |
| Mammalia (Mammals) | *Ornithorhynchus anatinus* | Platypus | TNKS1: XP_001508887.3<br>TNKS2: XP_028915823.1 |
| | *Sarcophilus harrisii* | Tasmanian devil | TNKS1: XP_031799023.1<br>TNKS2: XP_031811897.1 |
| | *Bos taurus* | Cattle | TNKS1: NP_001193089.1<br>TNKS2: XP_024841840.1 |
| | *Mus musculus* | House mouse | TNKS1: NP_780300.2<br>TNKS2: NP_001157107.1 |
| | *Homo sapiens* | Modern human | TNKS1: NP_003738.2<br>TNKS2: NP_079511.1 |

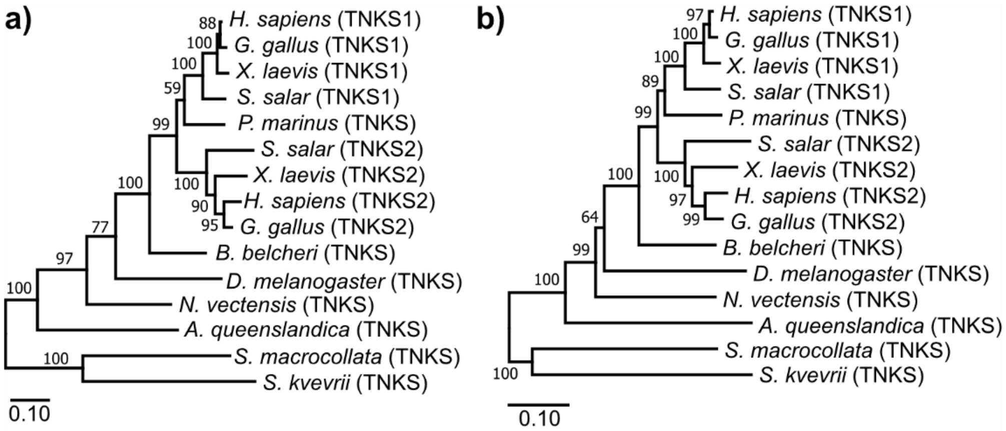


**Figure S1: Phylogenetic analysis of tankyrases; related to Figure 2.** Construction of the phylogenetic trees was done with Maximum Likelihood (a) or Neighbor Joining (b). Percentages for bootstrap support values are shown for each node. Bootstrap tests were done with 1000 bootstrap replicates. The analysis was performed with truncated sequences starting from the ARC3 domain to match the likely incomplete N-terminal sequence from *S. macrocollata*.

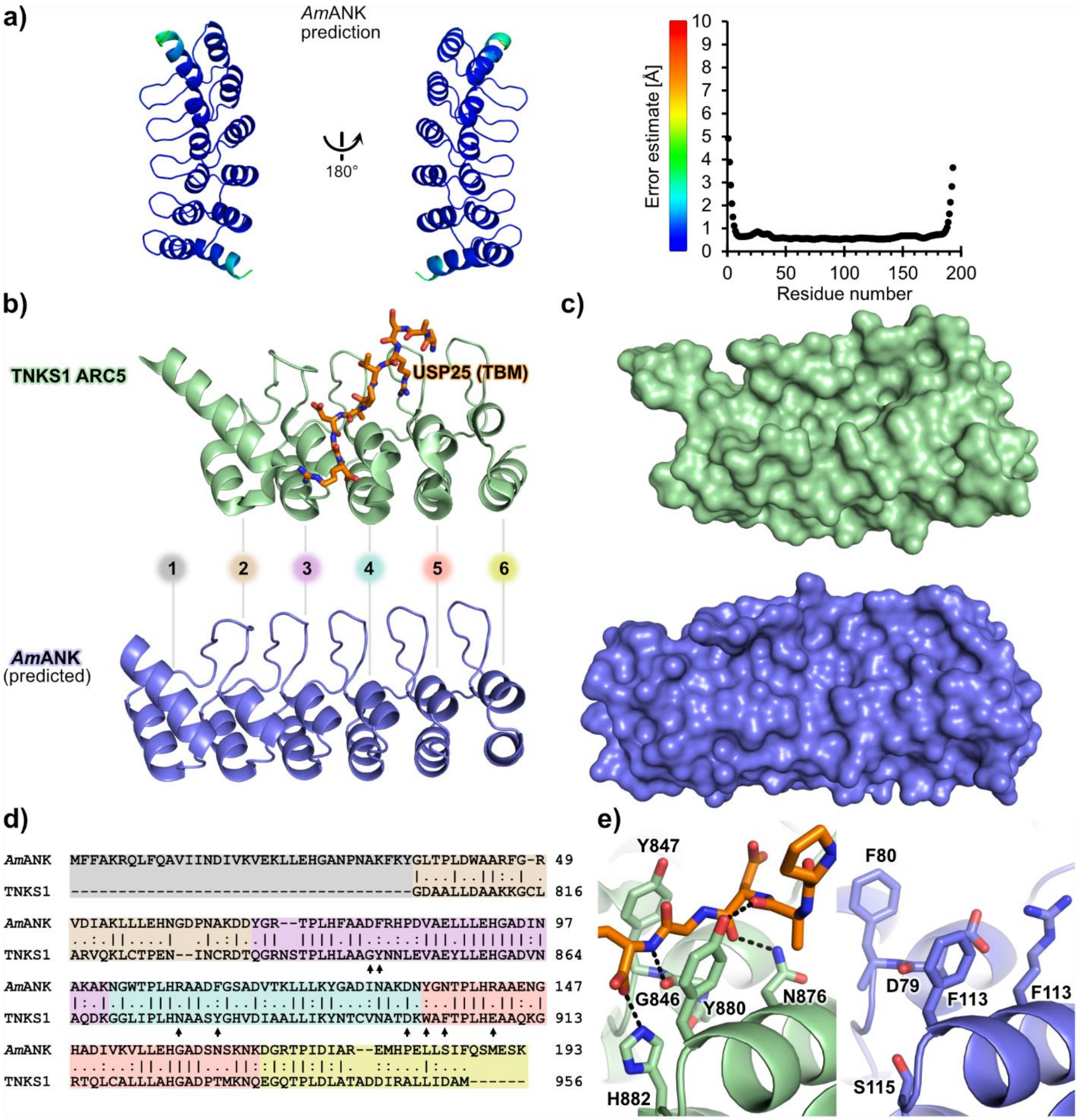


**Figure S2: Predicted structure of the TNKS-like ankyrin-repeat domain from the archaeon *Acidianus manzaensis*; related to Figure 2.** (a) The structure of the *A. manzaensis* ankyrin-repeat domain (*Am*ANK) was predicted by RoseTTAFold. Coloring of the model corresponds to the estimation of errors in Å. (b) Side-by-side view of the human TNKS1 ARC5 domain in complex with an USP25 TBM peptide (PDB: 5GP7) and the predicted structure of *Am*ANK. ARC5 contains 5 and *Am*ANK contains 6 ankyrin-repeat units. The ankyrin-repeat units are numbered. (c) Surface representations of TNKS1 ARC5 and *Am*ANK. While both structures are similar, the peptide-binding pocket in ARC5 appears more well-defined. (d) Pairwise sequence alignment of *Am*ANK and the ARC5 domain from TNKS1. The ankyrin-repeat units are highlighted in different colors. Important residues for TBM binding are indicated by arrows for TNKS1. (e) Close-up view of the aromatic glycine-sandwich sub-site in ARC5 binding the USP25 TBM peptide (left) and corresponding view of *Am*ANK (right). While the two tyrosines are replaced with phenylalanines in *Am*ANK, other residues facilitating this interaction differ and would likely not permit this mode of peptide-binding.

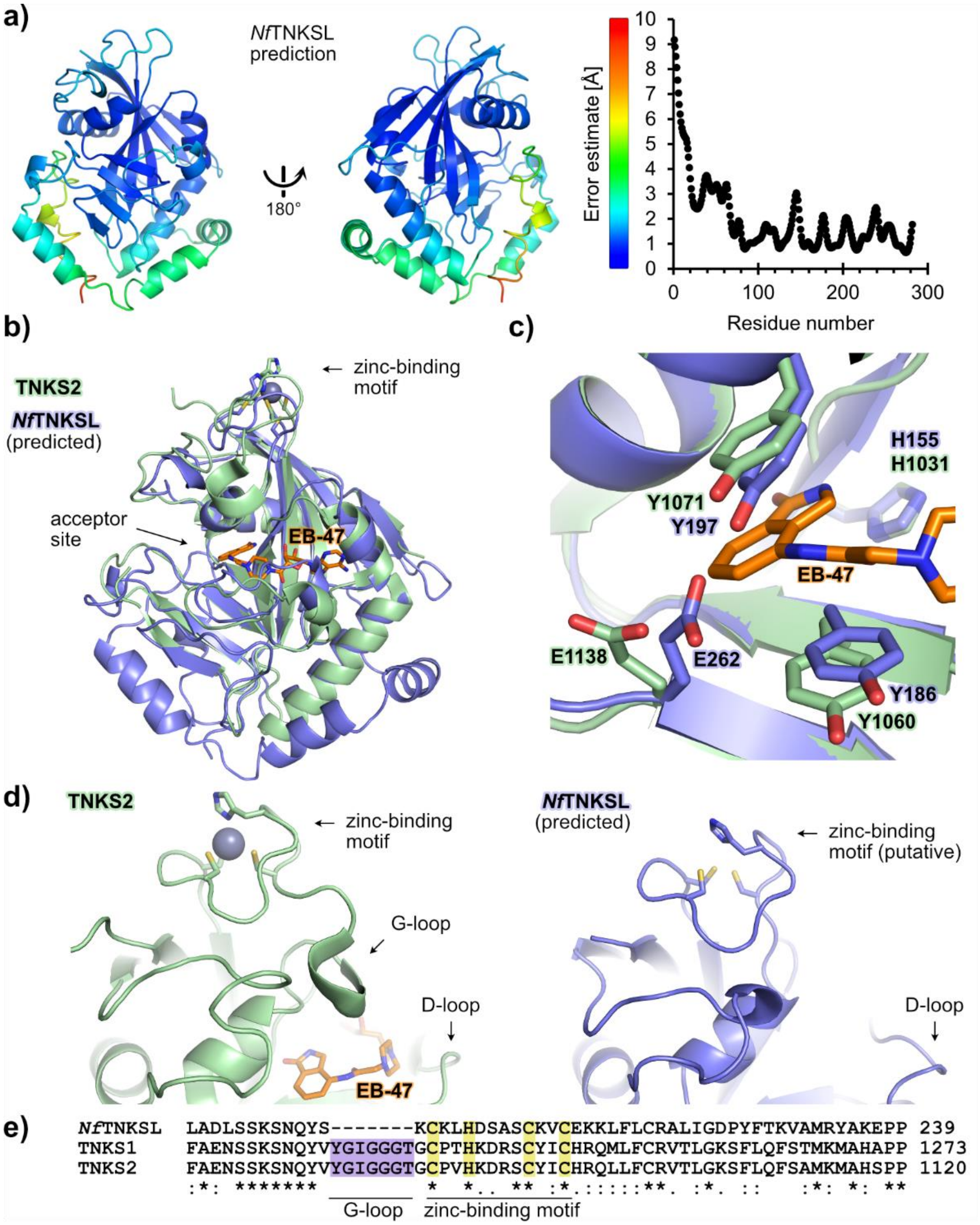


**Figure S3: Predicted structure of the TNKS-like ART domain from the amoeba *Naegleria fowleri*; related to Figure 2.** (a) The structure of the *N. fowleri* TNKS-like (*Nf*TNKSL) ART domain was predicted by RoseTTAFold. Coloring of the model corresponds to the estimation of errors in Å. (b) Superposition of the predicted *Nf*TNKSL structure and the crystal structure of TNKS2 ART domain in complex with the NAD$^+$-mimic EB-47 (PDB: 4BJ9). (c) Close-up view of the nicotinamide sub-pocket for the superimposed structures. (d) Side-by-side comparison of the structures from TNKS2 and *Nf*TNKSL. The unique zinc-binding motif in TNKSs was modelled in very similar conformation in the predicted *Nf*TNKSL structure, while the Glycine-rich loop (G-loop) present in TNKSs is absent in *Nf*TNKSL. (e) Multiple sequence alignment of ART domains from human TNKS1, human TNKS2 and *Nf*TNKSL. The G-loop region is colored in violet and not present in *Nf*TNKSL; the ligand residues from the zinc-binding motif are highlighted in yellow.

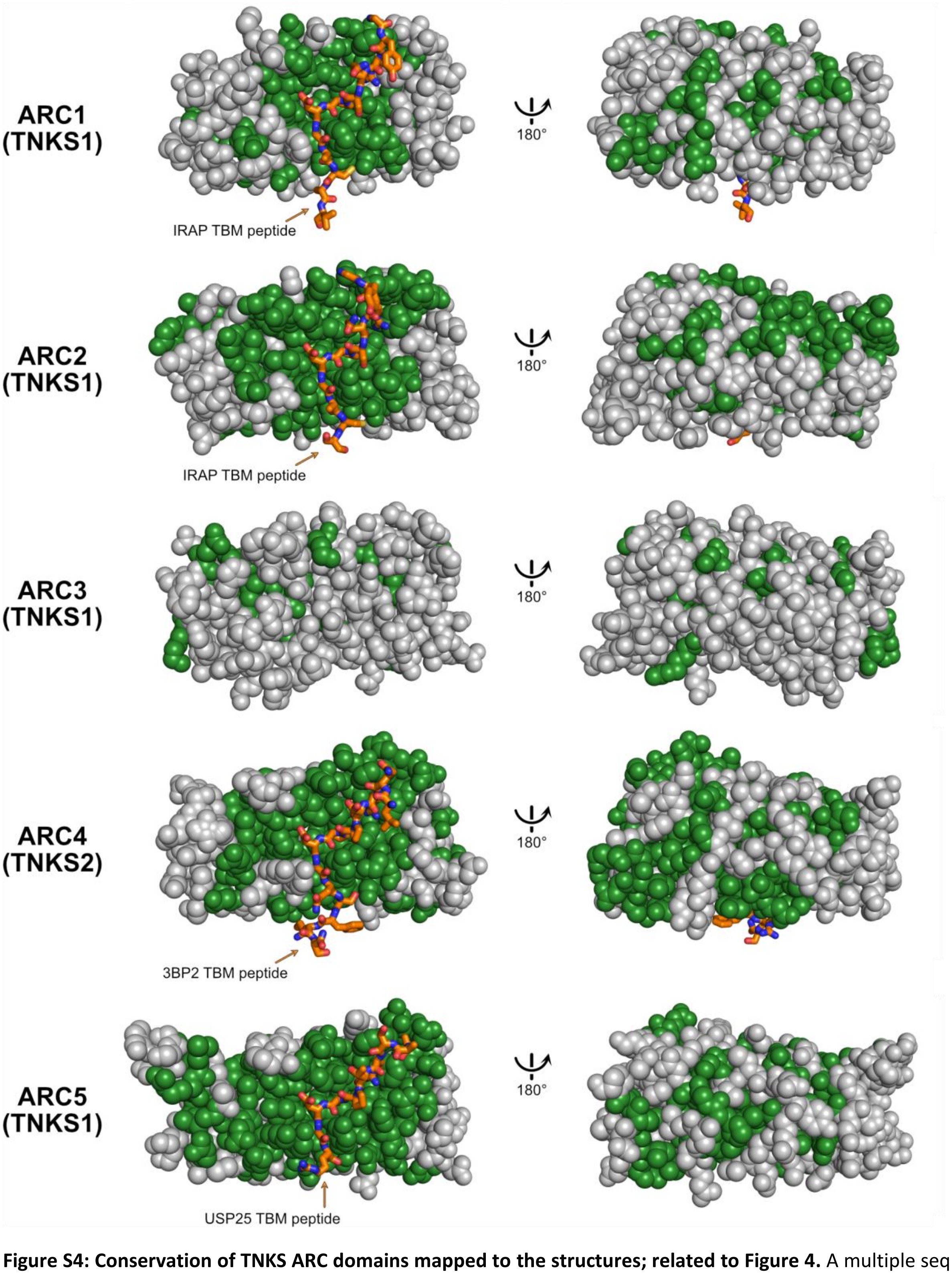


**Figure S4: Conservation of TNKS ARC domains mapped to the structures; related to Figure 4.** A multiple sequence alignment of TNKSs from 11 metazoan species (as exemplified in **Figure 4b**) was used to map all identical residues to the structures of TNKS ARC domains, shown in green: TNKS1 ARC1 (PDB: 5JHQ), TNKS1 ARC2 (PDB: 5JHQ), TNKS1 ARC3 (PDB: 5JHQ), TNKS2 ARC4 (PDB: 3TWR), TNKS1 ARC5 (PDB: 5GP7).

**a)**

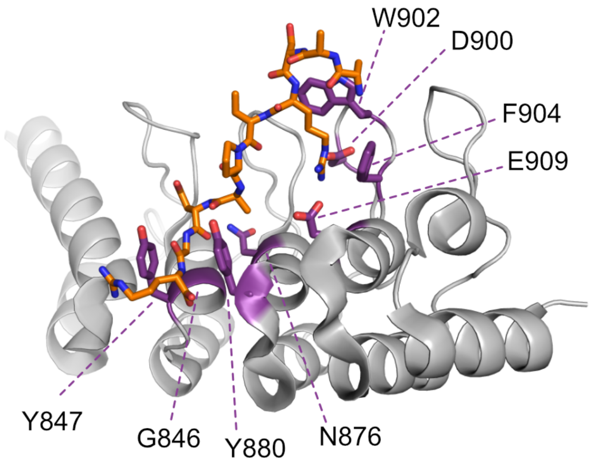


**b)**

```
                                                                  G846 Y847
TNKS1_ARC5       --GDAALLDAAKKGCLARVQKLCTPENINCRDTQGRNSTPLHLAA---GYNNLEVAEYLL
TNKS1_ARC3       EFKGHSLLQAAREADLAKVKKTLALEIINFKQPQSH-ETALHCAVASLHPKRKQVTELLL
PmTNKS_ARC3      EFRGHSLLQAAREADVTRVKKHLGLDIINFKHPQTH-ETALHCAVASPYPKRKQVSELLL
BbTNKS_ARC3      EFKGHCLLDGARQTDMAKVKKHLQLDIVNFKHPYTH-DTALHCAALSPYPKRRQITELLI
DmTNKS_ARC3      EYKGHCLLDACRKCDVSRAKKLVCAEIVNFVHPYTG-DTPLHLAVVSPDGKRKQLMELLT
AqTNKS_ARC3      EYRGYQLLAAAEDGGIILLKKLLSSQLLKFQHHQTL-DTLLHKAVLSKSSNRQSVIDALL
                    .  ** ....  :   :*    : ::  .     .* ** *.     :. .: : *

                                   N876 Y880           D900 W902 F904 E909
TNKS1_ARC5       EHGADVNAQDKGGLIPLHNAASYGHVDIAALLIKYNTCVNATDKWAFTPLHEAAQKGRTQ
TNKS1_ARC3       RKGANVNEKNKDFMTPLHVAAERAHNDVMEVLHKHGAKMNALDTLGQTALHRAALAGHLQ
PmTNKS_ARC3      RKGANVNEKTKDLLTPLHVASDKAHNDVIDVLLKHGAKINVTDSLGQTSLHRAAHAGHLT
BbTNKS_ARC3      RKGANLNDKNKEFLTPLHVASDKCYIDVMEVLLKHGAKVNALDSLGQTALHRGAHCGHVQ
DmTNKS_ARC3      RKGSLLNEKNKAFLTPLHLAAELLHYDAMEVLLKQGAKVNALDSLGQTPLHRCAR--DEQ
AqTNKS_ARC3      KRGINLNIGNKENMTPLICAAKKGILEVVEQLVQRGANINHQDINGMTSLHWAVQNEHAQ
                 .:*  :*   *  : **  *:.    :    * : .: :*  *  . * **  .

TNKS1_ARC5       LCALLLAHGADPTMKNQEGQTPLDLATADDIRALLIDAM
TNKS1_ARC3       TCRLLLSYGSDPSIISLQGFTAAQMGNEAVQQILS----
PmTNKS_ARC3      TCRLLLSHGCDPNLPSLQGFTAAQLGNESVQQILH----
BbTNKS_ARC3      ACRLLLSFGVDPSIVSLQGVTAAQMATEAVQQMLH----
DmTNKS_ARC3      AVRLLLSYAADTNIVSLEGLTAAQLASDSVLKLLK----
AqTNKS_ARC3      ICRYLLSSGANPSIVNNQGQTIYQLKTSDTIQLIL----
                     **: . : .: . :* *  :: .    : :
```

**Figure S5: The ARC3 domains in both human and early-diverging tankyrases are unlikely to bind TBM peptides.** (a) Residues important for binding TBMs by TNKS ARC domains are colored purple and mapped to the structure of ARC5 from human TNKS1 (PDB: 5GP7). The TBM peptide of USP25 is shown in orange. (b) Multiple sequence alignment of ARC5 from human TNKS1 and ARC3 sequences from human TNKS1, *Petromyzon marinus* TNKS (*Pm*TNKS), *Branchiostoma belcheri* TNKS (*Bb*TNKS), *Drosophila melanogaster* TNKS (*Dm*TNKS) and *Amphimedon queenslandica* TNKS (*Am*TNKS). Residues important for binding TBMs are highlighted in purple for TNKS1 ARC5.

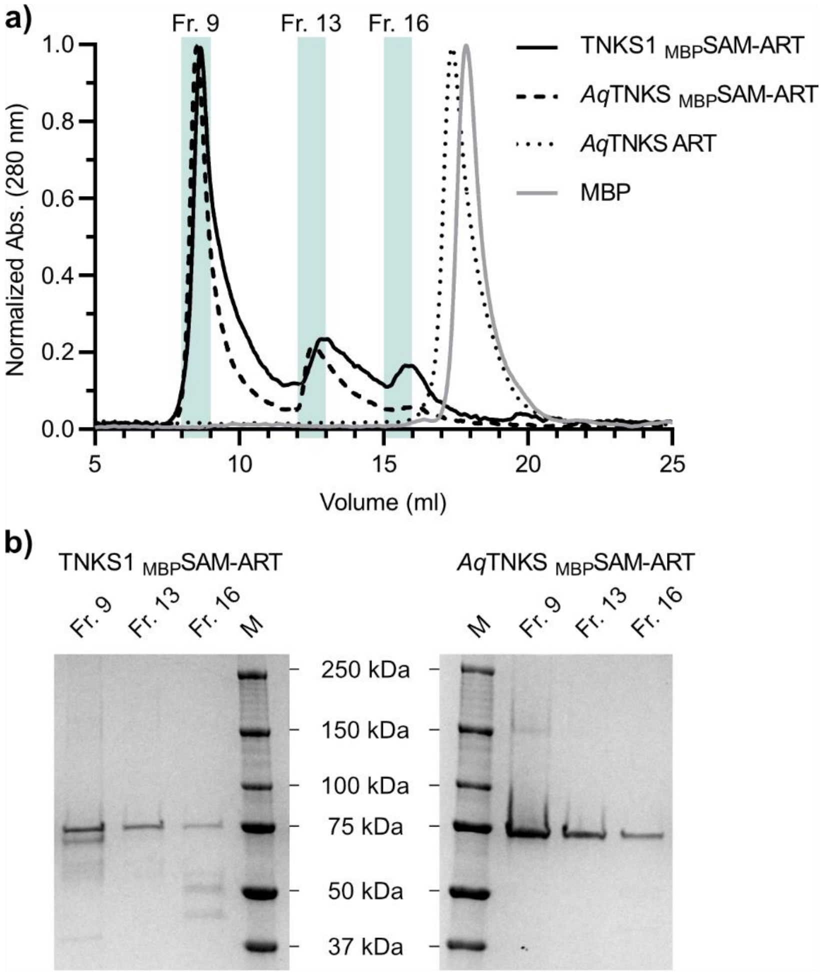


**Figure S6: Size-exclusion chromatography of TNKS1 and *Aq*TNKS constructs; related to Figure 5.** a) Size-exclusion chromatogram of TNKS constructs or MBP. In separate runs, 0.5 mg of each protein was loaded to a Superdex 200 Increase 10/300 GL column at a flowrate of 0.5 ml/min at 4°C. The absorbance at 280 nm was monitored and fractions (Fr.) of 1 ml were taken. (b) For the SAM-ART constructs of TNKS1 or *Aq*TNKS, fractions 9, 13 and 16 were analyzed by SDS-PAGE. Precision Plus Protein All Blue (BioRad) was used as protein weight marker (M).
